# Using null models to compare bacterial and microeukaryotic metacommunity assembly under shifting environmental conditions

**DOI:** 10.1101/752303

**Authors:** Máté Vass, Anna J. Székely, Eva S. Lindström, Silke Langenheder

## Abstract

Temporal variations in microbial metacommunity structure and assembly processes in response to shifts in environmental conditions are poorly understood. Hence, we conducted a temporal field study by sampling rock pools in four-day intervals during a 5-week period that included strong changes in environmental conditions due to intensive rain. We characterized bacterial and microeukaryote communities by 16S and 18S rRNA gene sequencing, respectively. Using a suite of null-model approaches to assess dynamics in community assembly, we found that strong changes in environmental conditions induced small but significant temporal changes in assembly processes and triggered different responses in bacterial and microeukaryotic metacommunities, promoting distinct selection processes. Incidence-based approaches showed that the assemblies of both communities were mainly governed by stochastic processes. In contrast, abundance-based methods indicated the dominance of historical contingency and unmeasured factors in case of bacteria and microeukaryotes, respectively, which we distinguished from dispersal-related processes using additional tests. Taken together, our study highlights that community assembly processes are not static, and the relative importance of different assembly processes can vary under different conditions and between different microbial groups.

## Introduction

Different assembly processes such as environmental selection, dispersal and/or stochastic processes can simultaneously influence community composition [1]. The relative importance of the different processes is highly context–dependent and dynamic and may therefore vary in importance over time [2–5] as a consequence of processes such as ecological succession [4, 6], seasonality [3, 5, 7] or changes in connectivity between sites [8, 9]. Despite the increased recognition that community assembly processes are not static, the majority of studies is based on snapshot sampling which cannot adequately capture their dynamics [7].

Besides contemporary changes in environmental conditions and dispersal processes [10], past environmental conditions and dispersal events (i.e., historical contingency) may also influence the temporal dynamics of assembly processes [11–13]. For instance, changes in the variation in environmental heterogeneity could affect the relative importance of species-sorting or selection processes [8], while changes in dispersal rates could affect the possibilities for mass effects [14] or the extent of dispersal limitation [15]. Further, the importance of historical contingency may also depend on the environmental context. For example, priority effects – the impact of particular species on community development due to prior arrival at a site – may be affected by environmental disturbances that initiate colonization events that intensify the importance of the phenomenon [11]. Several studies detected a trajectory from stochastic to deterministic assembly processes in time following a disturbance [16, 17], which might reflect effects of initially strong, but transient priority effects that diminish over time as more species arrive and establish in the post-disturbance community. Finally, the probability of priority effects may also increase when productivity is high [18], because the growth of early colonizers is promoted [11].

Only a few studies have directly compared assembly mechanisms between different groups of microorganisms such as bacteria and microeukaryotes. Based on the differences in e.g., cell sizes, generation times and life history traits, differences in assembly processes are expected between these two groups [19–21]. For instance, it has been suggested that marine protist communities are governed by species-sorting to a greater extent than are marine bacterial communities [22, 23], while on the contrary, other studies indicated the opposite [21]. Microeukaryotes have been suggested to be mainly shaped by stochastic mechanisms (i.e. drift) [21, 24] and to be more subject to the effect of dispersal than bacteria [25]. Hence, there are to date conflicting results on how assembly processes differ and persist through time in bacterial and microeukaryotic communities.

The statistical ‘toolbox’ that is currently used to gain insights into the importance of different community assembly processes consists of several complementary approaches that all have their own strengths and limitations. Recently, null model approaches that quantitatively compare assembly processes have been increasingly used [26, 27]. For example, the elements of metacommunity structure (EMS) method allows to distinguish randomly assembled communities from those assembled by species-sorting processes [28, 29]. The incidence-based (Raup-Crick) beta-diversity (β_RC_) [30] has been used to differentiate between deterministic and stochastic assembly processes [18, 31]. In addition, based on the assumption that phylogenetic relatedness is indicative of shared environmental response traits [32], null model approaches have been extended to integrate phylogenetic information [33]. Specifically, Stegen et al. [15, 27] have combined null model approaches based on phylogenetic and abundance-based (Raup-Crick) beta-diversity (β_RCbray_) measures to quantitatively estimate the relative importance of processes such as selection, drift, dispersal limitation and mass effects. Furthermore, this quantitative process estimate (QPE) method also differentiates between heterogenous/variable (i.e., beta-diversity enhancing) and homogeneous (i.e., beta-diversity diminishing) selection processes.

We carried out an extensive field study of bacterial and microeukaryotic communities in rock pools, which are particularly variable habitats both in space and time. The above-mentioned statistical approaches were applied to assess the temporal changes in community assembly processes. We hypothesized that temporal changes in the importance of different assembly processes should occur in dependence on changes in environmental conditions and, further, that these changes differ between bacterial and microeukaryotic communities.

## Material and methods

### Sampling procedure

Samples were taken from 20 neighboring rock pools – referred to as a ‘metacommunity’ – located along the Baltic Sea coast on the island of Gräsö, Sweden (60°29’54.0” N, 18°25’48.9” E) (Supplemental Fig. S1). The rock pools were sampled ten times, starting on 14 August 2015 and ending on 19 September 2015 in four-day intervals (Fig. S2). Four of the pools dried out at certain occasions during the sampling period. Intensive rain (starting August 31) occured in the middle of the study period and separated a cooler wet period (air temperature (°C): 13.98±1.35, precipitation (mm): 3.97±6.94, wind speed (m/s): 6.7±2.91) from an extended dry period (air temperature (°C): 17.71±0.98, precipitation (mm): 0.09±0.36, wind speed (m/s): 5.6±1.61) prior to the rain (Fig. S2).

Numerous abiotic and biotic variables were measured at each sampling occasion. Specifically, conductivity and temperature were measured using a WTW Conductometer (Cond 3210 SET 2 incl. TetraCon 325-3 measuring cell, Germany). Morphological parameters such as maximum length, width and depth were recorded for each pool. Zooplankton samples were collected by filtering 2 l of water through a net (250 μm) and fixed with 70% ethanol for subsequent analysis. Five litres of water were collected in sterile plastic bottles and transported back to the laboratory where the samples were further processed. For quantification of bacterial abundance, samples were preserved with sterile-filtered formaldehyde at a final concentration of 2% and stored at 4 °C, while for bacterial and eukaryotic community analyses, pre-filtered (250 μm) water samples (100–500 ml) were collected by vacuum filtration onto 0.2 μm 47 mm membrane filters (Supor-200, Pall Corporation, Port Washington, NY, USA), and then, stored at –80 °C until further processing (see below).

### Sample analysis

Nutrient concentrations, such as total phosphorus (TP) and total nitrogen (TN) were measured spectrophotometrically (Perkin Elmer, Lambda 40, UV/VIS Spectrometer, Massachusetts, USA) and by catalytic thermal decomposition method (Shimadzu TNM-L, Kyoto, Japan), respectively according to standard procedures. Water colour was determined by measuring the absorbance of GF/C-filtered (Whatman® Glass microfiber filter, Sigma-Aldrich, USA) water at 436 nm. Chlorophyll-a was measured [34] as an estimator of the biomass of primary producers. Bacterial abundance was determined by flow cytometry as in Székely et al. [35] with the modification of using 2.27 μM of SYTO 13 fluorescent nucleic acid stain (Invitrogen, Eugene, Oregon, USA).

For both bacterial and micro-eukaryotic community composition analyses, DNA extraction was performed from the membrane filters (PowerSoil DNA Isolation Kit, MoBio Laboratories Inc., Carlsbad, CA, USA). Bacterial 16S rRNA and eukaryotic 18S rRNA genes were amplified with bacterial (341F and 805R; [36]) and eukaryotic (574*f and 1132r; [37]) primer constructs, respectively. A full step-by-step protocol for the detailed two-step PCR protocol has been deposited in the protocols.io repository (dx.doi.org/10.17504/protocols.io.468gzhw). Amplicons were sequenced at the SciLifeLab SNP&SEQ Technology Platform (host by Uppsala University) using Illumina MiSeq v3 sequencing chemistry. The raw sequencing data are available at the European Nucleotide Archive under accession number PRJEB30954. A detailed report of the data processing is provided in the supplementary material. The taxonomic composition of both datasets is visualized in Supplemental Figure S3 and S4.

### Statistical analysis

All statistical analyses (see Fig. 1 for an overview) and visualizations were conducted in R version 3.3.2 [38].

**Figure 1.**
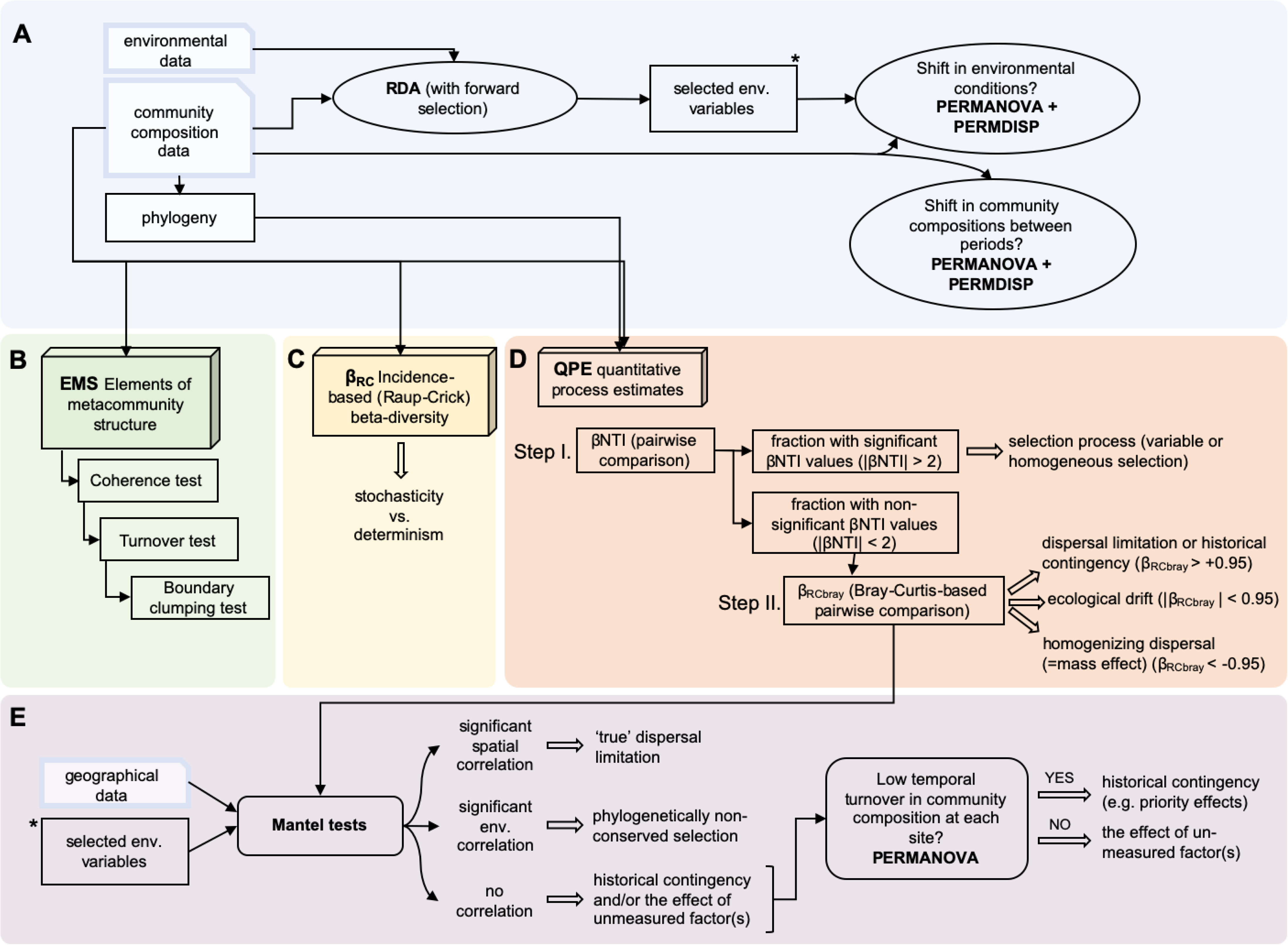
Flow chart of the statistical analyses performed in this study. (A) Redundancy analysis with forward selection was performed to select the most important environmental variables that explain variation in the community matrices. Then, we compared the variance and homogeneity of environmental and community distances between the dry and wet period using PERMANOVA and PERMDISP, respectively. (B-C-D) Three null-model approaches were applied. (B) EMS identifies metacommunity properties emerging in site-by-OTUs incidence matrix [28, 29]. (C) Incidence-based (Raup-Crick) beta-diversity (β_RC_) tests stochasticity and determinism using a metric provided by Chase et al. [31]. (D) QPE quantifies assembly processes involving phylogeny and abundance-based (Raup-Crick) beta-diversity (β_RCbray_) following the framework of Stegen et al. [27]. (E) We performed (partial) Mantel tests as a complement to the QPE between β_RCbray_ and geographical and environmental distance matrices in order make a clear distinction of historical contingencies (e.g. priority effects) and/or unmeasured factors, phylogenetically non-conserved selection and pure effects of dispersal limitation. Then, we distinguished historical contingency and the effects of unmeasured factors by assessing temporal change of community composition at the level of individual rock pool using PERMANOVA.

#### Shifts in environmental conditions structuring communities

We excluded the four rock pools that occasionally dried out from all analyses. Then, we accounted for collinearity among standardized environmental variables by omitting highly collinear variables (Pearson |r| > 0.7) based on Dormann et al. [39]. To select the variables most strongly associated with the variance of the observed communities, we applied redundancy analysis (RDA) on the Hellinger-transformed sequence data with forward selection (based on 999 permutations; variables retained at *p* < 0.05), separately for bacteria and microeukaryotes (Fig. 1A) (see Results section for details). Differences in means and variances in selected variables between the dry and wet periods were tested using Kruskal-Wallis test and Levene’s test, respectively. Additionally, to assess the separation between the two periods, permutational multivariate analysis of variance (PERMANOVA) with 999 permutations was performed using the function ‘adonis’, further, the multivariate homogeneity of group dispersions (PERMDISP) was tested using the function ‘betadisper’ in the package ‘vegan’ [40].

#### Elements of metacommunity structure (EMS)

Elements of metacommunity structure (EMS) were assessed for each time point following the frameworks developed by Leibold & Mikkelson [28] and Presley et al. [29] (Fig. 1B). EMS enables to identify metacommunity properties that emerge in a site-by-species incidence matrix that is compared with null model expectations obtained through randomization [41]. Random matrices were produced by the ‘r1’ method (fixed-proportional null model). For this, the matrices (OTU table from 16S and 18S rRNA gene sequences, separately) were ordinated according to the primary axis via reciprocal averaging and then hierarchically analysed using three tests (coherence, turnover and boundary clumping) (for more details, see supplementary material). The package ‘metacom’ [41] was used to detect any pattern of metacommunity structure related to an idealized scenario (‘metacommunity type’). Following the suggested hierarchical framework of EMS, we specifically focused on the outcome of coherence tests (the number of embedded absences in the ordinated matrix and comparing the empirical value to a null distribution) in the subsequent statistical analyses since the majority of metacommunities were associated with checkerboard and random metacommunity types. Differences in the coherence (z-values) between the two periods (wet and dry) were tested using a Kruskal-Wallis test.

#### Incidence-based beta-diversity (β_RC_)

The incidence-based (Raup-Crick) dissimilarity indices (β_RC_) were calculated to test whether community were stochastically or deterministically assembled (Fig. 1C). For this, we used the ‘raup_crick’ function provided by Chase et al. [31]. When β_RC_ is not significantly different from 0, the community is considered to be stochastically assembled. β_RC_ values close to −1 indicate that communities are deterministically assembled and more similar to each other than expected by chance, whereas β_RC_ values close to +1 indicate that deterministic processes favor dissimilar communities. The averaged dissimilarities for each time point and for each dataset (bacteria and microeukaryotes) were calculated separately. Differences in β_RC_ between the two periods (wet and dry) were tested using a Kruskal-Wallis test.

#### Quantitative process estimates (QPE)

To quantify the relative importance of potential species sorting, dispersal limitation, drift and mass effects (we refer to this as ‘quantiative process estimates – QPE’ throughout the manuscript), we followed the two-step framework developed by Stegen et al. [27] (Fig. 1D). This approach requires that phylogenetic distances (PD) among taxa reflect differences in the ecological niches they inhabit, thus, carry a phylogenetic signal. The presence of phylogenetic signals was tested using Mantel correlograms, as described in Stegen et al. [27]. We found that niche differences caused by most of the environmental variables that structured communities according to the RDA results (see above) could induce turnover in phylogenetic community composition (Fig. S5, S6) and thereby fulfill the prerequisite of this framework.

To perform QPE based on pairwise comparisons, firstly, we determined to what extent the observed βMNTD (β-mean-nearest-taxon-distance) deviated from the mean of the null distribution and evaluated significance using the β-Nearest Taxon Index (βNTI; difference between observed βMNTD and the mean of the null distribution in units of SDs). If the observed βMNTD value is significantly greater (βNTI > 2) or less (βNTI < −2) than the null expectation, the community is assembled by variable or homogeneous selection, respectively. If there is no significant deviation from the null expectation, the observed differences in phylogenetic community composition should be the result of dispersal limitation, homogenizing dispersal (mass effect) or random drift. To estimate the relative importance of these processes, in the second step, the abundance-based (Raup-Crick) beta-diversity was calculated using pairwise Bray-Curtis dissimilarity (β_RCbray_) [27]. Based on the calculated β_RCbray_, we can assume that communities that were not selected in the first step, thus not assembled by selection, were structured by (i) dispersal limitation coupled with drift if β_RCbray_ > +0.95, (ii) homogenizing dispersal if β_RCbray_ < −0.95, or (iii) random processes acting alone (drift) if β_RCbray_ falls in between −0.95 and +0.95 (Fig. 1D). The first fraction, β_RCbray_ > +0.95, may either indicate ‘true’ effects of dispersal limitation and/or history contingency that both result in more dissimilar communities than expected by chance. Hence, throughout the manuscript we use the term ‘dispersal limitation or historical contingency’ for this fraction. Differences in the QPE between the two periods (wet and dry) were tested using Kruskal-Wallis test.

To further assess whether ‘true’ dispersal limitation might have occurred, Mantel correlations between bacterial and microeukaryotic community dissimilarities (β_RCbray_) and geographic distances (Euclidean distances of geographical coordinates) were done using Pearson correlation and 999 permutations (Fig. 1E). By this, we could confirm any potential dispersal limitation and reject historical contingency if there is significant correlation between β_RCbray_ and spatial distance. Further, Mantel correlation analyses between β_RCbray_ and environmental dissimilarities (Euclidean distances) were done to test if community dissimilarities were possibly due to selection by environmental factors that lack a phylogenetic signal (‘phylogenetically non-conserved selection’) and was therefore not detected in the first step but instead was retained in the second step of the QPE analysis. To determine if significant geographic distance effects were confounded by effects of spatially autocorrelated environmental variation and vice versa, partial Mantel tests were used with the respective third matrix as covariate for time points where both correlations with geographic and environmental distances were significant.

## Results

### Temporal variation in relevant environmental variables

RDA with forward selection showed that conductivity (F = 11.37, *p* = 0.005), water temperature (F = 2.99, *p* = 0.005), nutrients (TP: F = 4.71, *p* = 0.005 and TN: F = 2.04, *p* = 0.005), depth (F = 1.62, *p* = 0.03) and Daphnia abundance (F = 1.62, *p* = 0.015) correlated significantly with the variation in bacterial community composition. For microeukaryotes, the same variables and copepod abundance were significant (conductivity: F = 8.34, *p* = 0.005; water temperature: F = 4.05, *p* = 0.005; TN: F = 2.84, *p* = 0.005; TP: F = 1.99, *p* = 0.005; depth: F = 2.68, *p* = 0.005; Daphnia abundance: F = 1.87, *p* = 0.015; copepod abundance: F = 2.04, *p* = 0.005).

The temporal fluctuations of the selected variables followed similar patterns during the sampling period (Fig. 2). Specifically, we found that there was a clear separation point in the middle of the study period (31 August, between two sampling occasions on 30 August and 3 September; dashed line in Fig. 2) from when on environmental conditions became more homogenous, i.e., the variance across the rock pools decreased (except in the case of depths and conductivity, although the latter one was marginally insignificant) (Table S1). These differences supported our separation of the study period and corresponding datasets into two periods, a dry and wet period (Figs 2 and S2). Consequently, the pools had higher mean water temperature, conductivity, zooplankton abundance, and nutrient concentrations, lower mean pool depth and more spatially heterogeneous conditions (high variance across pools) in the dry compared to the wet period (Fig. 2, Table S1). This separation was further supported by PERMANOVA and PERMDISP analyses, which showed that the environmental conditions (F = 31.07, *p* = 0.001; Fig. S7) and their variances (F = 79.58, *p* < 0.001) clearly differed between the two periods. There were also significant differences in the composition of the bacterial and microeukaryotic communities of the dry and wet period but no difference in their homogeneity (beta-dispersion) (Figs S8, S9). Meanwhile, at the level of individual pools significant differences in community composition (PERMANOVA) were detected in some of the pools (7 out of 16) for bacteria and for most pools (15 out of 16) in the case of microeukaryotes (Table S2) without any difference in their beta-dispersion (PERMDISP, Table S3).

**Figure 2.**
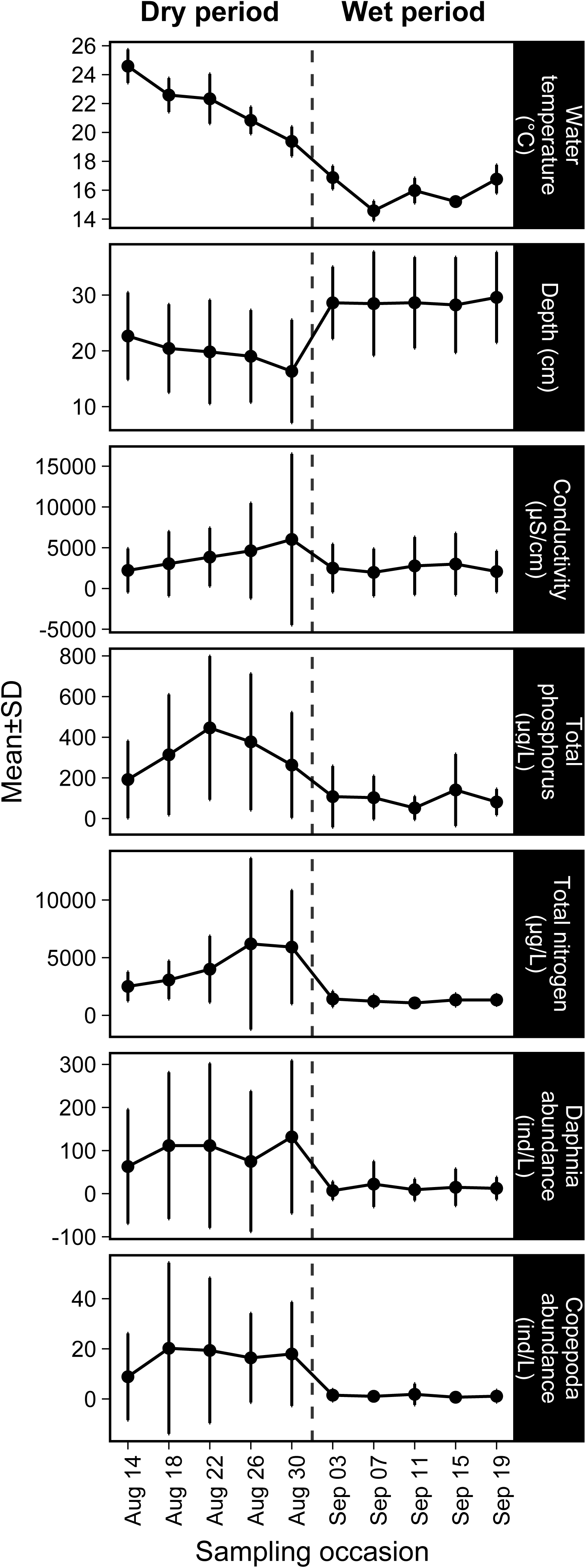
Temporal dynamics of the mean values and standard deviations of environmental variables that significantly affected either the composition of bacterial or the microeukaryotic communities (based on RDA) during the study period. The dashed line indicates rain that separated the study period into a dry and wet period.

### Elements of metacommunity structure (EMS)

In general, the observed z-value of coherence did not show wide variation across the bacterial and microeukaryotic datasets, which were shaped by random processes at the majority of the time points in both cases. Checkerboard pattern emerged at one occasion and two occasions for the bacterial and microeukaryotic metacommunities, respectively, while a nested, clumped species loss pattern was detected once during the wet period in bacteria (Fig. 3, Table S4). Microeukaryotic metacommunities were also mainly characterized by random patterns, except for two occasions when checkerboard patterns occurred (Fig. 3). There was no significant change of coherence (z-values) over time in any of the observed datasets (bacteria: χ_dry vs. wet_ = 0.884, *p* = 0.347; microeukaryotes: χ_dry vs. wet_ = 3.153, *p* = 0.076), however, there were trends towards slightly higher coherence (z-values) in the wet period compared to the dry period, especially in the microeukaryote communities.

**Figure 3.**
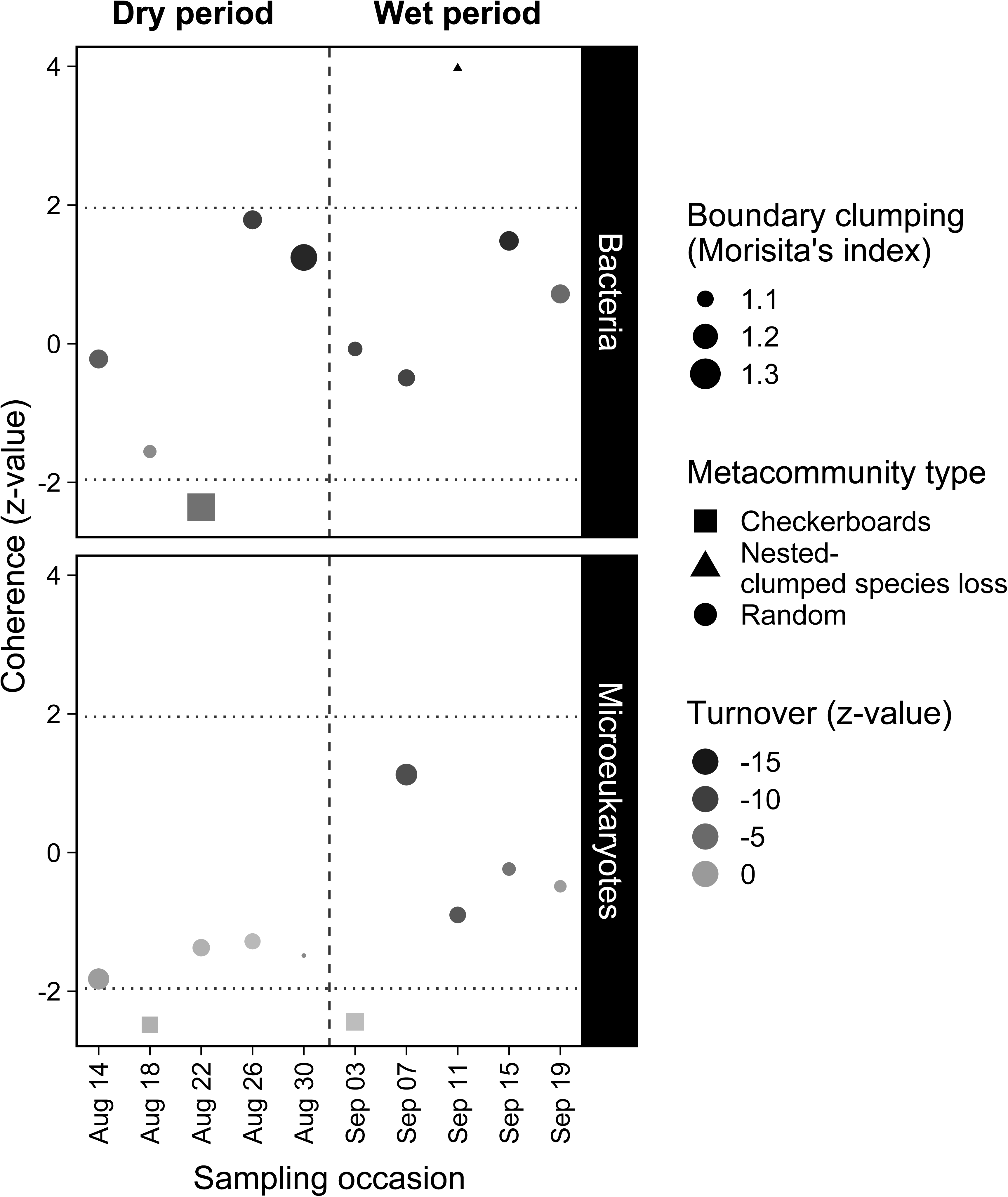
Temporal variation of metacommunity types of the bacterial and microeukaryotic datasets. Within the dotted lines (−1.96 < coherence z-value < 1.96) metacommunities are randomly structured. Positive significant values (coherence z-value > 1.96) indicate that species’ distribution occur in response to environmental variation. Significantly negative coherence (coherence z-value < −1.96) indicates checkerboard distribution. The greyscale represents species turnover (z-value; number of observed replacements compared to a null distribution) where positive values indicate species replacements in response to environmental variation and negative values nested species distributions caused by species losses. The size of the symbols denotes the Morisita’s index (boundary clumping) which shows the degree of spatial distribution of species in a metacommunity where lower numbers indicate over-dispersed boundaries and higher numbers clumped boundaries. Vertical dashed lines refer to the division between the dry and wet period.

#### Incidence-based beta-diversity (β*_RC_*)

Across the 16 rock pools the average β_RC_ varied within a narrow range, not deviating strongly from the null expectations (0.066–0.227 and −0.221–0.197 in bacterial and microeukaryotic communities, respectively), which indicates stochastic assembly. For bacteria, there was no clear pattern or trend in β_RC_ associated with the shift in the environmental conditions (χ_dry vs. wet_ = 0.273, *p* = 0.602) (Fig. 4). For microeukaryotes the β_RC_ values decreased at the beginning of the wet-period, but thereafter they increased rapidly (Fig. 4), although they remained within the ‘stochastic’ range (−0.95 < β_RC_ < +0.95) (χ_dry vs. wet_ = 0.535, *p* = 0.465).

**Figure 4.**
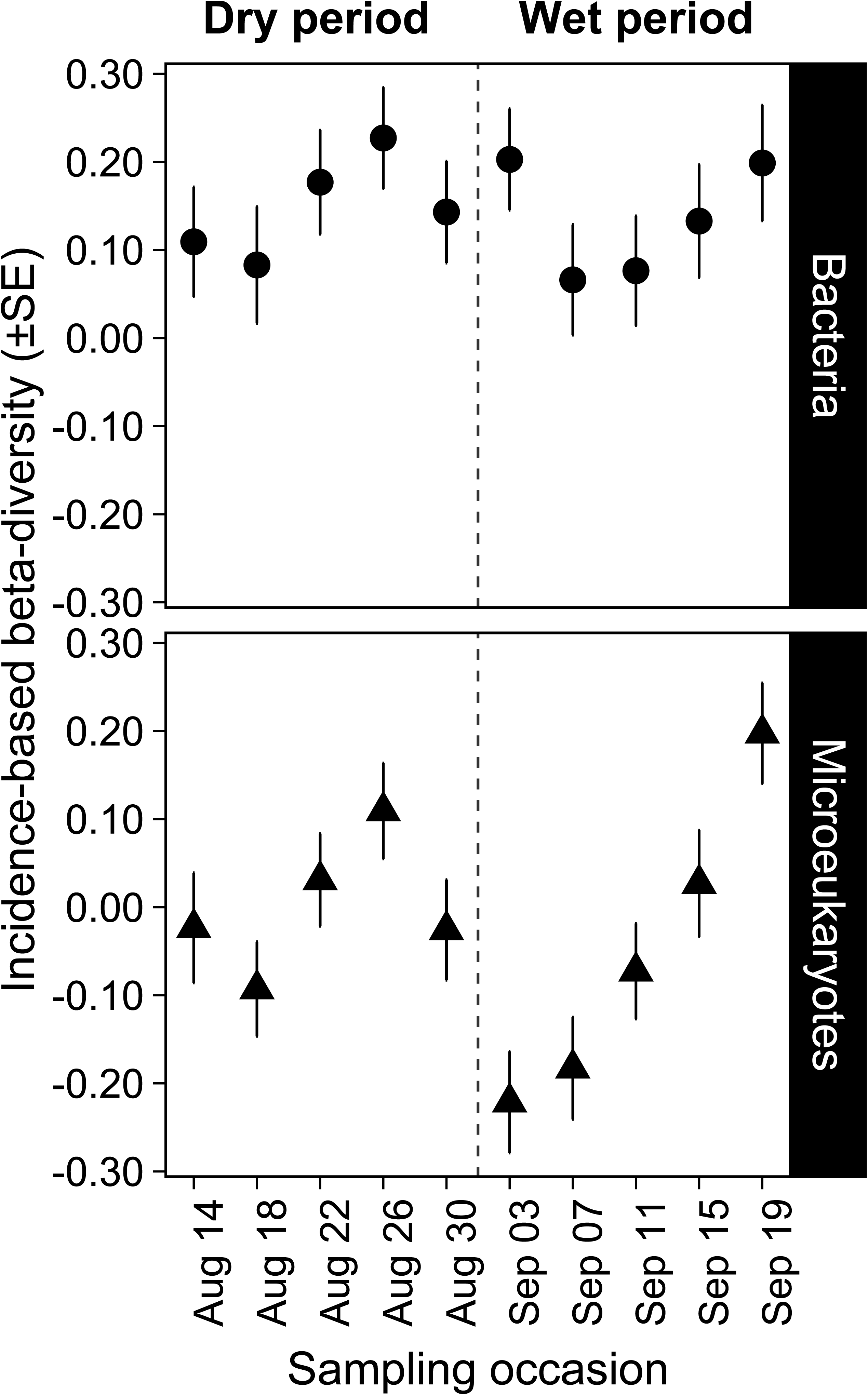
Variation of incidence-based (Raup-Crick) beta-diversity (β_RC_) for bacteria and microeukaryotic communities during the study period. Dashed line refers to the division between dry and wet period.

### Quantitative process estimates (QPE)

The quantitative process estimates showed temporal variation over the study period with some differences between the two organism groups (Fig. 5). For bacteria, dispersal limitation or historical contingency was the dominant assembly processes (60.95–80.83% of all pairwise comparisons) followed by homogeneous selection processes (4.17–27.62%), random processes (drift, 4.76–15.38%), variable selection (0–12.5%) and homogenizing dispersal (0–1.67%). The relative proportion of homogeneous selection increased in the wet period (χ_dry vs. wet_ = 5.34, *p* = 0.021), while that of variable selection decreased compared to the dry period, although this decline was not significant (χ_dry vs. wet_ = 1.32, *p* = 0.251). There were no significant changes detected between the two periods in the case of dispersal limitation or historical contingency (χ_dry vs. wet_ = 1.10, *p* = 0.293), drift (χ_dry vs. wet_ = 0.01, *p* = 0.916) and homogenizing dispersal (χ_dry vs. wet_ = 0.05, *p* = 0.828). For microeukaryotic metacommunities, dispersal limitation or historical contingency was also the dominating assembly process at all time points (56.19–85.83%). The second and third most dominant assembly processes were drift (5.00–22.86%) and variable selection (1.67–18.1%), respectively, whereas the proportions of homogeneous selection (0–2.86%) and homogenizing dispersal (0–2.56%) were negligible. The proportion of dispersal limitation or historical contingency decreased (χ_dry vs. wet_ = 5.77, *p* = 0.016) while variable selection increased during the wet period after the first rainfall (χ_dry vs. wet_ = 4.81, *p* = 0.028), whereas the slight increase of drift during the wet-period was not significant (χ_dry vs. wet_ = 2.81, *p* = 0.094). The importance of homogenizing dispersal differed between the two periods (χ_dry vs. wet_ = 4.51, *p* = 0.034), while opposite to the bacterial metacommunities, homogeneous selection did not change significantly (χ_dry vs. wet_ = 0.41, *p* = 0.522) (Fig. 5).

**Figure 5.**
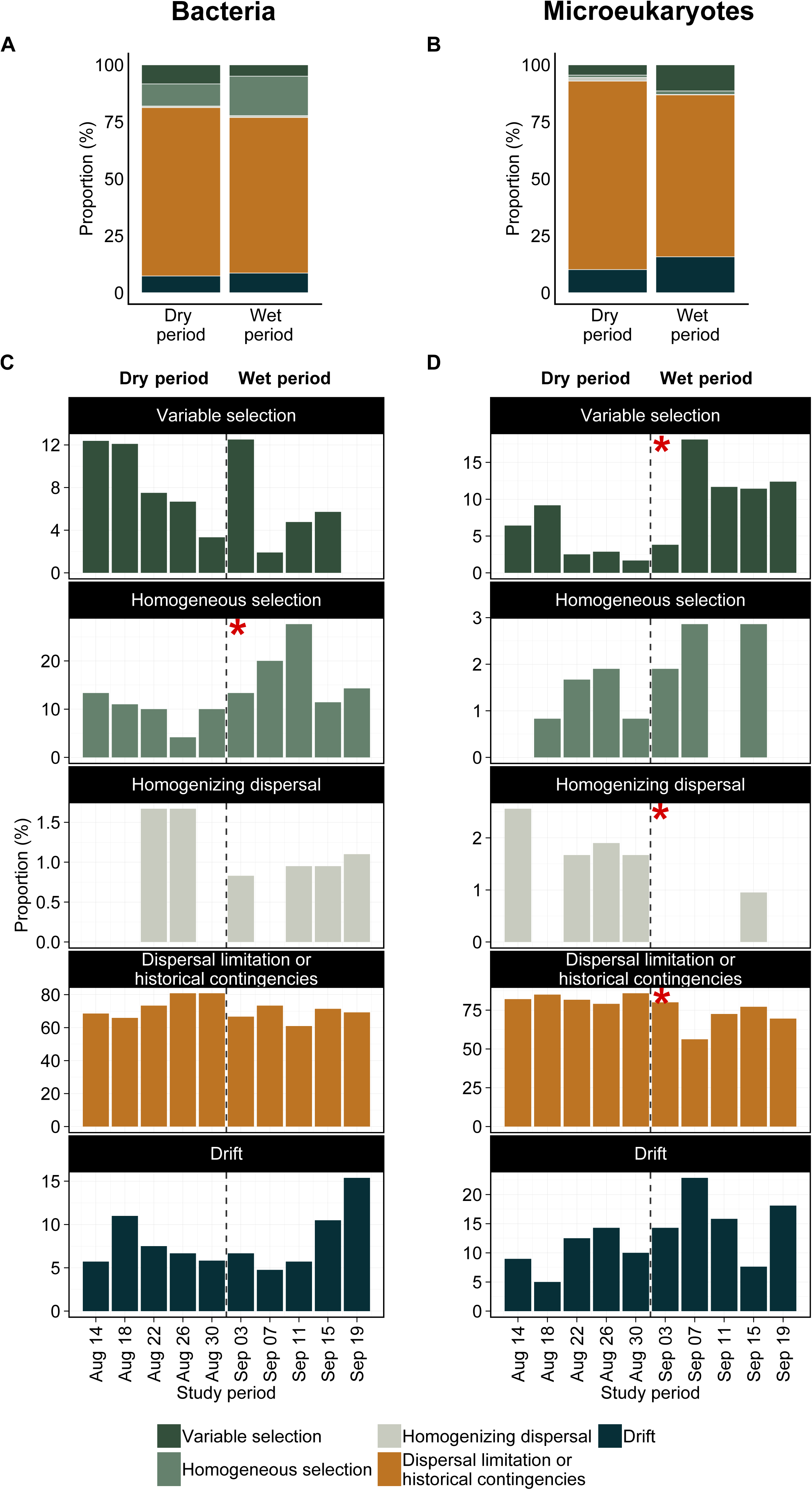
Overall (A, B) and temporal (C, D) dynamics of the relative importance of different community assembly processes expressed as the proportion of community pairs assembled either by species-sorting (variable or homogeneous selection), dispersal limitation or historical contingency, homogenizing dispersal or drift for bacteria (A, C) and microeukaryotic (B, D) communities. Note that the scales are not equal on the C and D facet plot. The dashed lines refer to the division between the dry and wet period, and red asterisks indicate significant differences between them (Kruskal-Wallis test, significance at *p* < 0.05 level).

Mantel correlations between community distance matrices (βRCbray, the fraction retrived for the second step of QPE) and geographical/environmental distance matrices were generally weak, showed no consistent pattern, and were only significant for a few time points (Fig. 6). Microeukaryotic communities showed significant correlations for geographic distance in one case and for both environmental and geographic distances in another. In the latter case the correlations were even significant when controlled for effects of covariation by environmental distance in cases of geographic distance (partial r_M_ = 0.23, *p* = 0.003) or geographic distance in case of environmental distance (partial r_M_ = 0.19, *p* = 0.032), respectively. Meanwhile, bacterial community compositions were significantly correlated to environmental distance only once.

**Figure 6.**
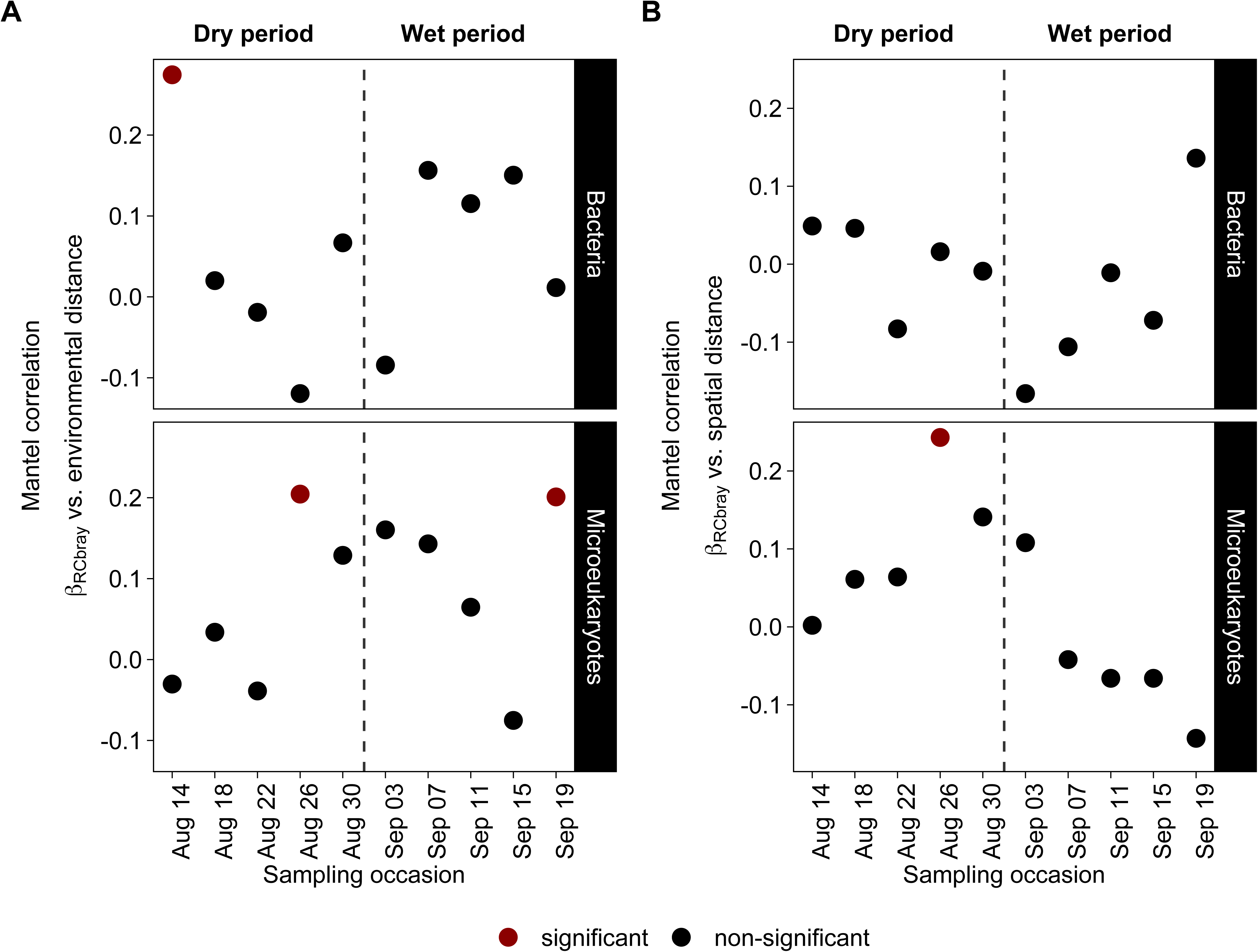
Mantel (Pearson) correlations between bacterial and microeukaryotic community dissimilarities (abundance-based Raup-Crick beta-diversity – β_RCbray_) and (A) environmental and (B) geographic distances (Euclidean distances) for each sampling occasion. The dashed lines refer to the division between dry and wet period.

## Discussion

### Environmental dependency of assembly mechanisms

Intensive rain from the middle of our rock pool sampling campaign separated the study period into a distinct dry and wet period, allowing us to specifically investigate the environmental dependency of assembly mechanisms of microbial communities over time. According to the incidence-based beta-diversity (β_RC_) patterns, both bacterial and microeukaryotic communities were primarily stochastically assembled throughout the study period (β_RC_ ≈ 0) despite the observed environmental changes during the transition from the dry to the wet period. Testing the elements of metacommunity structure (EMS) provided further evidence for stochastic assembly. These results resonate the idea that microbial community assembly is unpredictable because stochastic occurrence patterns are due to rapid population dynamics [42]. However, in contrast to the results from the β_RC_ and EMS analyses, the quantitative process estimates (QPE) framework showed that both bacterial and microeukaryotic community assemblies were dominated by dispersal limitation or historical contingency at all time points. The relative importance of dispersal limitation or historical contingency in microeukaryotes was significantly higher in the dry period compared to the wet period, while in bacterial communities it increased towards the end of dry period, but then decreased slightly during the wet-period (Fig. 5). This suggests that a lack of connectivity among pools during the dry period lead to a temporary enforcement of dispersal limitation or history contingency (see discussion below for our interpretation). Further, the environmental shift between the dry and wet period slightly promoted the influence of homogeneous selection and variable selection for bacteria and microeukaryotes, respectively, even though none of the changes in environmental conditions that occurred throughout the study period induced strong selection processes.

While our study gives overall support for the dominance of stochastic and dispersal limitation or historical processes in the assembly of microbial communities, previous studies of bacterial communities in rock pools have shown that either both selection processes (i.e. species sorting) alone [43] or both environmental and spatial effects shape the communities [44]. However, it has also been shown that the importance of environmental vs. spatial effects over time varies in response to changes in environmental conditions [5]. Here, we provide a more refined picture of temporal changes in assembly processes that occur at much shorter temporal scales compared to the previous studies. Since community assembly is dynamic our temporal study provides a more comprehensive understanding of how microbial communities respond to environmental changes on short-time scales compared to previous snapshot studies [43, 44] or a study where changes were analysed over longer time periods as well as longer sampling intervals [5]. The present study also differs from the previous ones in that a broader suit of statistical methods was applied, allowing the analysis of further assembly mechanisms than in the studies where primarily variation partitioning was used [5, 45].

### Comparison of null model approaches

Our results show that different null model approaches led to different conclusions about the dominant community assembly processes. Generally, the key differences among the applied three null model approaches are that EMS and β_RC_ are developed for detecting patterns in binary presence-absence matrices based on taxonomic beta-diversity estimates only, while the QPE framework is based on quantitative, abundance-based matrices integrating phylogenetic information. One possible explanation why selection processes were only detected by the QPE but not the non-quantitative methods (EMS and β_RC_) could be that species sorting is to a great extent related to changes in the relative abundances of species but not the replacement of species. This highlights that abundance-based metrics might be more suited to describe microbial beta-diversity and the underlying assembly mechanisms at spatial scales similar to those studied here [46, 47].

### Differences between bacterial and microeukaryotic communities

Based on the results of null model approaches the hypothesis that bacteria and micro-eukaryotes are assembled by different assembly processes was partly supported. More specifically, the relative importance of assembly processes, and the way they changed in response to changing environmental conditions differed for bacterial and microeukaryotic communities. More specifically, for the bacterial metacommunity, there was a slight decrease in the influence of variable selection processes in the wet period compared to the previous dry period, while the relative importance of homogeneous selection processes significantly increased, which is conform with the idea that homogenization in environmental conditions among rock pools leads to more similarly composed bacterial communities [5]. On the contrary, for microeukaryotes, homogeneous selection processes remained negligible throughout the study period period, while the relative influence of variable selection surprisingly increased in the wet period. One possible explanation might be that in the wet period the increased water depth could have generated more gradients within each pool for environmental parameters such light, which is crucial for photo- and mixotrophic microeukaryotes [48, 49], thus, promoting the establishment of distinct local microeukaryotic communities. In general, it is worth to mention that both the bacterial and the microeukaryotic dataset consisted of several distinct groups of organisms which have very different population dynamics, niche-preferences and interspecific interactions (Fig. S5, S6). This could potentially mask important selection forces that act at each taxonomic level. More specifically, when a metacommunity consists of sets of species that are more structured by environment and others that are less so, a comprehensive perspective (pooling all groups together) could result in a fuzzy, stochastic picture of assembly [50]. Hence, a separate investigation of different microeukaryotic and bacterial groups (e.g. heterotrophs vs. autotrophs) might reveal different influences of assembly mechanisms [3].

### Historical contingency vs. pure dispersal limitation

As mentioned earlier the quantitative process estimate analysis showed that both bacterial and microeukaryotic communities were primarily shaped by dispersal limitation or historical contingencies. At a first glance, the dominance of dispersal limitation seems surprising, given the idea that has persisted in microbiology for a long time that microorganisms are to a great extent not dispersal limited. This idea has now been challenged as many studies have, for instance, detected spatial distance effects for microorganism [5, 44, 51, 52]. Moreover, other studies using quantitative process estimates have also shown considerable proportion of ‘dispersal limitation or historical contingency’ fraction [22, 53]. However, problems related to the interpretation of dispersal limitation fraction have also been raised [54], because it does not purely reflect dispersal limitation but rather a number of different processes, such as historial contingency and effects of phylogenetically non-conserved selection processes. To more explicitly test whether pure dispersal limitation was present in our study, we used Mantel correlations of abundance-based Raup-Crick beta-diversity (β_RCbray_, on the fraction retrived for the second step of QPE) vs. geographical distances between pools to detect spatial distance-decay relationships [55]. However, except for one case in microeukaryotes, this was not the case neither for bacteria nor for microeukaryotes (Fig 7) and we do therefore not have robust support for dispersal limitation. Likewise, there was also no indication that the dispersal limitation or historical contingency fraction masked substantial effects of phylogenetically non-conserved selection processes related to measured environmental factors as Mantel correlations between RC_bray_ and environmental distance were also not significant in most cases. Therefore, it seems most likely that the dispersal limitation or historical contingency fraction points to the importance of the outcome of historical contingency and the effect of unmeasured factors (e.g. light) that are not phylogenetically conserved. Evidence for historical contingency, such as priority effects, would be a low temporal turnover in community composition at the level of individual rock pools despite the drastic environmental shift that occurred during the study period. In case of bacteria, most of the individual rock pools (9 out of 16) did not experience significant compositional shifts between the two periods. Hence, this suggests that these nine communities might have been influenced by priority effects, while the remaining pools might have been influenced by unmeasured environmental factors that are phylogenetically non-conserved. In contrast, microeukaryotic communities are unlikely to have experienced priority effects because most (15 out of 16) of the individual pools experienced significant compositional shift between the two periods (Table S2). Still, we could not explain these compositional shifts by spatial or measured environmental factors using Mantel tests (Fig. 6), suggesting that unmeasured environmental factors, such as light or trophic interactions are more important for microeukaryotes compared to bacteria. In summary caution needs to be taken when interpreting the results of quantitative process estimates and future refined statistical frameworks should integrate additional analyses such as those presented here to provide a more clear distinction of historical contingencies (e.g. priority effects), phylogenetically non-conserved selection and pure effects of dispersal limitation.

### Conclusions

Our results show that historical contingency and selection processes can play a key role in shaping microbial communities, but that the relative contribution of selection processes varies depending on the temporal variation of environmental heterogeneity and between bacteria and microeukaryotes. Furthermore, this present study highlighted that incidence-based and abundance-based null model approaches lead to different conclusions about the dominant community assembly process in microbial communities. Further, the outcomes of the current QPE framework act merely as a guide, because the fraction expected to indicate dispersal limitation may in reality depict other processes such as historical contingency, phylogenetically non-conserved selection, or even other, unmeasured processes. Our findings also show that temporal observations with high-resolution can provide more a comprehensive understanding than snapshot studies. Taken together, we encourage future studies to consider temporal variation of metacommunities and its environmental dependency, regardless the microbial group of interest, as well as to consider historical contingency (e.g. imprints of assembly history) as a potentially important assembly process.

## Acknowledgements

We thank Christoffer Bergvall for laboratory guidance, Christian Stolpe for laboratory assistance, and David Spange for creation of the site map, furthermore, Ron Coleman, Javier Vargas Calle and Jan Johansson for their help during field sampling. This study was funded by the Swedish Research Council. Further financial support was received by grants to A.J.S from the Swedish Research Council Formas and a Marie Curie International Outgoing Fellowship within the 7th European Community Framework Programme.

