## Supplementary material for "Using null models to compare bacterial and microeukaryotic metacommunity assembly under shifting environmental conditions"

**Supplementary Data and Methods for:** Using null models to compare bacterial and microeukaryotic metacommunity assembly under shifting environmental conditions

Máté Vass*, Anna J. Székely, Eva S. Lindström, Silke Langenheder

Department of Ecology and Genetics/Limnology, Uppsala University, Sweden

*Sampling site*

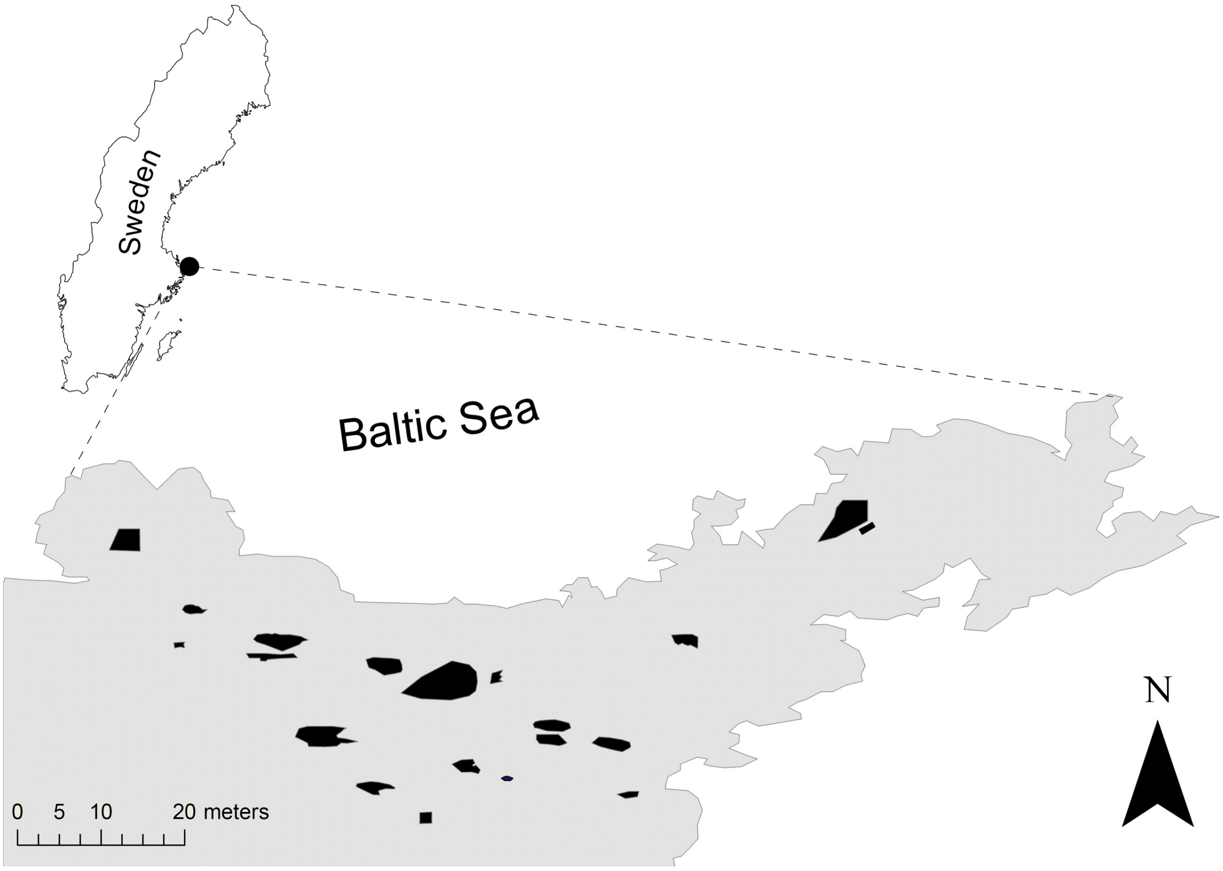


**Figure S1.** Sampled rock pools along the Baltic Sea coast on the island of Gräsö, Sweden

*Meteorological conditions*

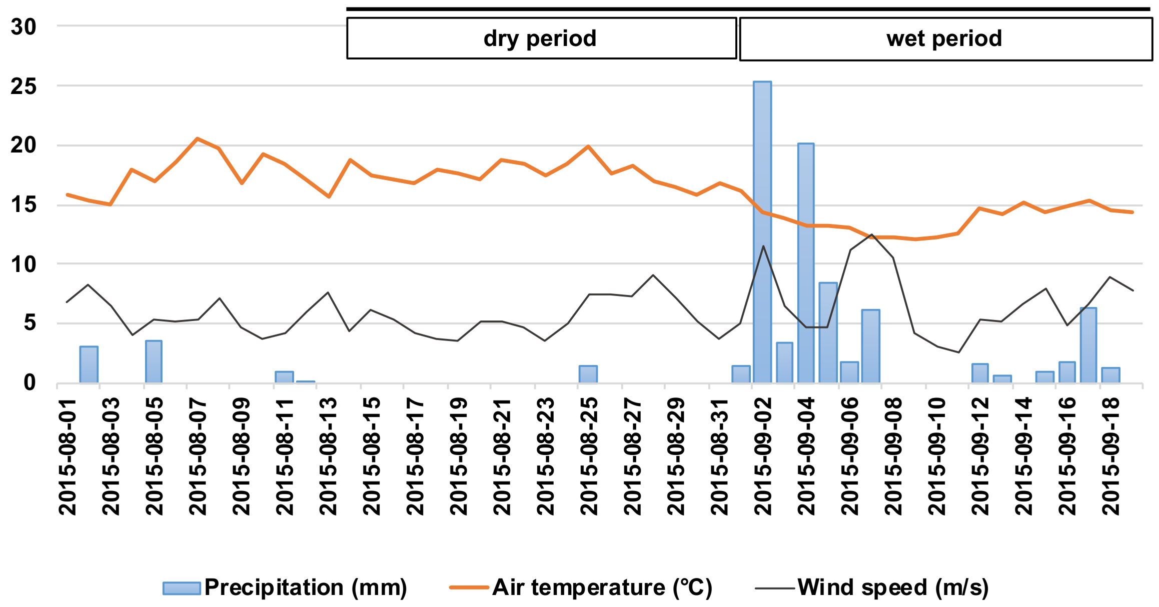


**Figure S2.** Daily total precipitation (mm), daily mean air temperature (°C) and daily mean wind speed (m/s) conditions at the Örskär meteorological station obtained from the Swedish Meteorological and Hydrological Institute (SMHI). The black bar refers to the study period.

*Taxonomic classification and distribution of OTUs found in the rock pools*

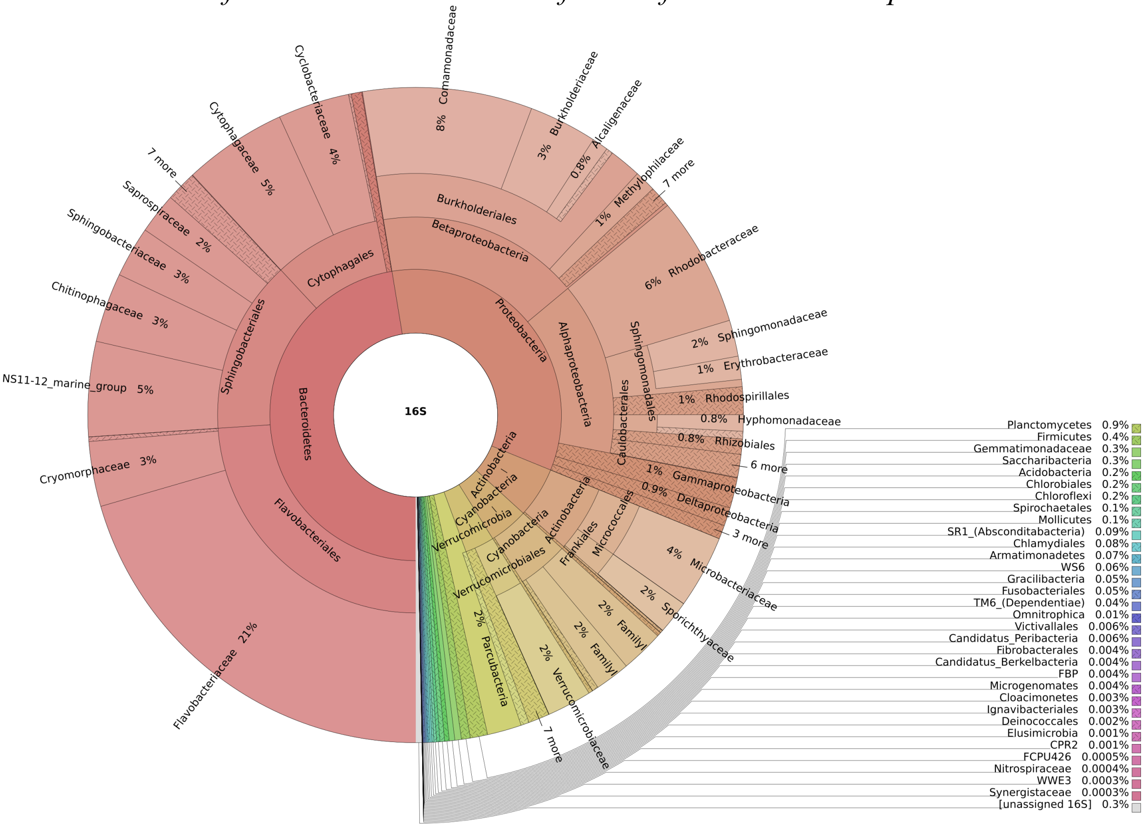


**Figure S3.** Taxonomic classification and distribution of bacterial OTUs (total of 4,587 OTUs)

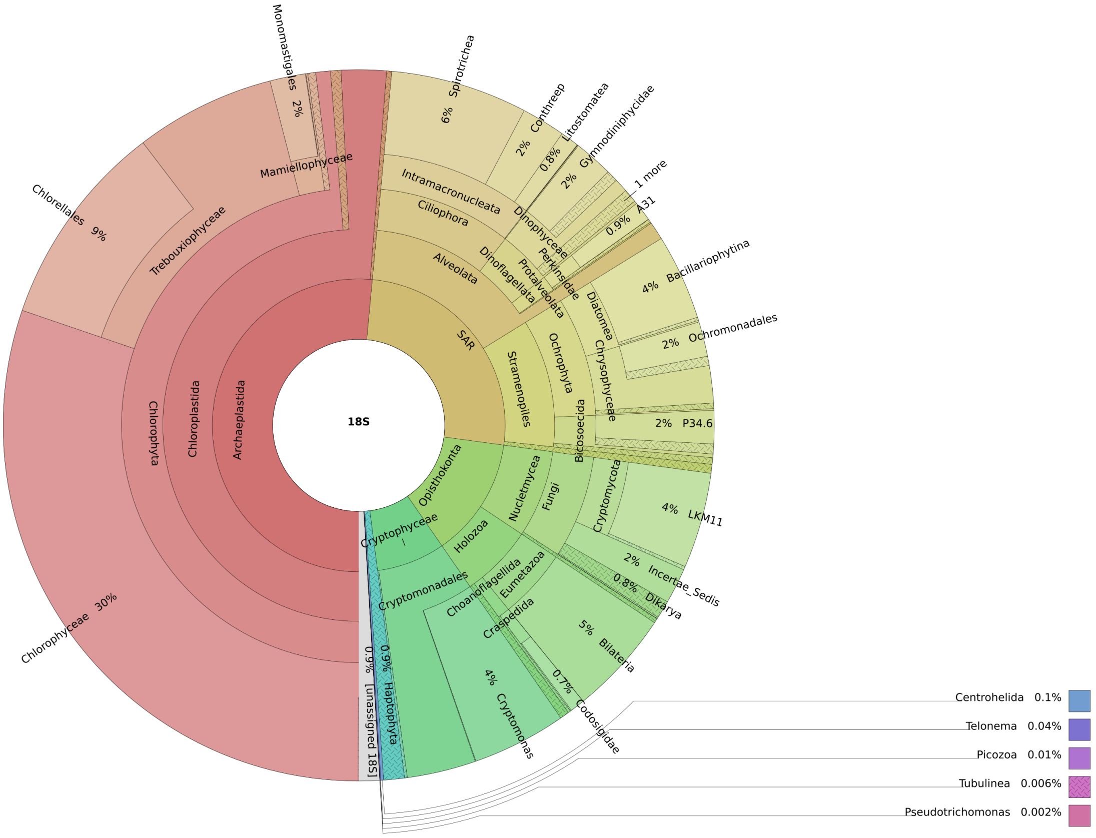


**Figure S4.** Taxonomic classification and distribution of microeukaryotic OTUs (total of 1,336 OTUs)

*Phylogenetic signals*

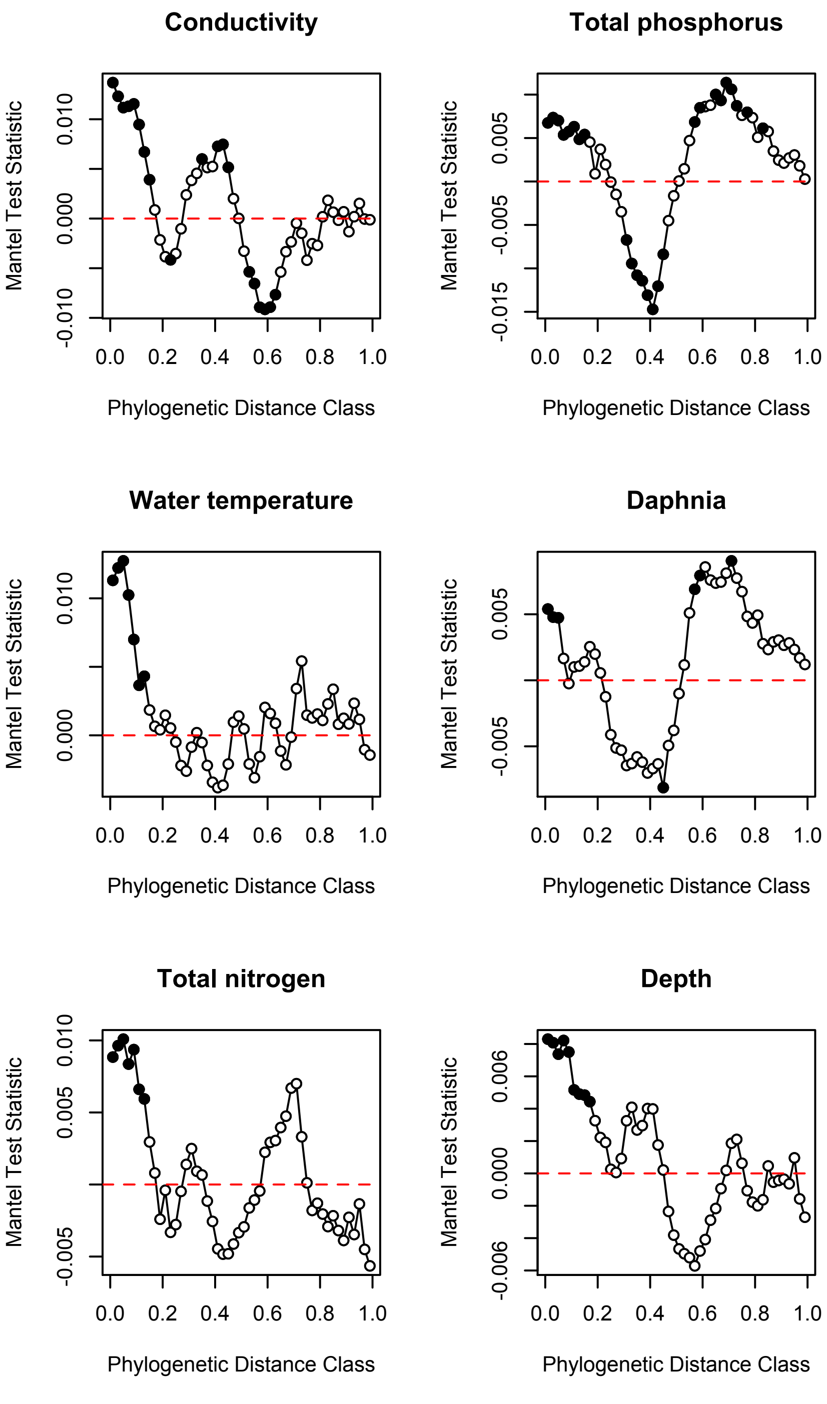


**Figure S5.** Phylogenetic Mantel correlograms showing significant phylogenetic signals across short phylogenetic distances (PD) in bacterioplankton communities. Closed dots denote significant correlations, relating between-OTU niche differences to between-OTU PDs across a given PD. Estimates of optimal OTU environmental niches were calculated for conductivity, total phosphorus, water temperature, Daphnia abundance, total nitrogen and rock pool depth.

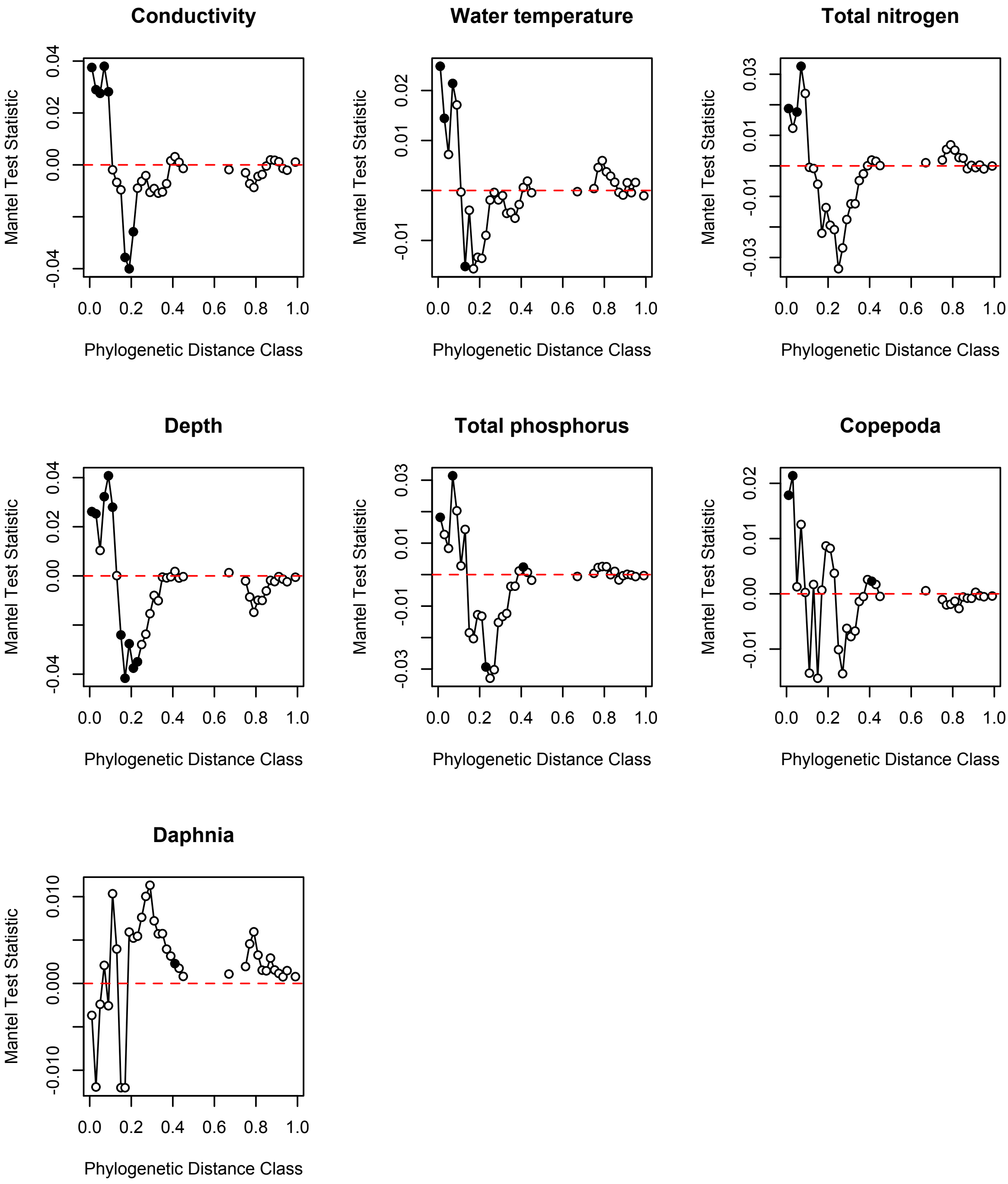


**Figure S6.** Phylogenetic Mantel correlograms showing significant phylogenetic signals across short phylogenetic distances (PD) in microeukaryotic communities (except in case of Daphnia abundance). Closed dots denote significant correlations, relating between-OTU niche differences to between-OTU PDs across a given PD. Estimates of optimal OTU environmental niches were calculated for conductivity, water temperature, total nitrogen, rock pool depth, total phosphorus, Copepoda and Daphnia abundances.

*Evidences for shifting environmental conditions in the rock pools during the study period and its effect on bacterial and microeukaryotic community compositions*

**Table S1.** Characteristic differences of rock pools between the dry and wet period. A Kruskal-Wallis test was used to compare the means, while Levene's test was used to compare the homogeneity of variance for each environmental variable. Significant values ($p < 0.05$) are in bold.

| | **Mean** | | **Variance** | |
|---|---|---|---|---|
| | *chi-squared* | *P* | *F* | *P* |
| Conductivity | 4.972 | **0.0257** | 3.586 | 0.059 |
| Copepod abundance | 31.525 | **<0.0001** | 44.68 | **<0.0001** |
| Daphnia abundance | 5.536 | **0.0186** | 24.43 | **<0.0001** |
| Depth | 49.047 | **<0.0001** | 0.335 | 0.563 |
| Total nitrogen | 109.4 | **<0.0001** | 16.319 | **<0.0001** |
| Total phosphorus | 70.002 | **<0.0001** | 24.96 | **<0.0001** |
| Water temperature | 143.09 | **<0.0001** | 32.42 | **<0.0001** |

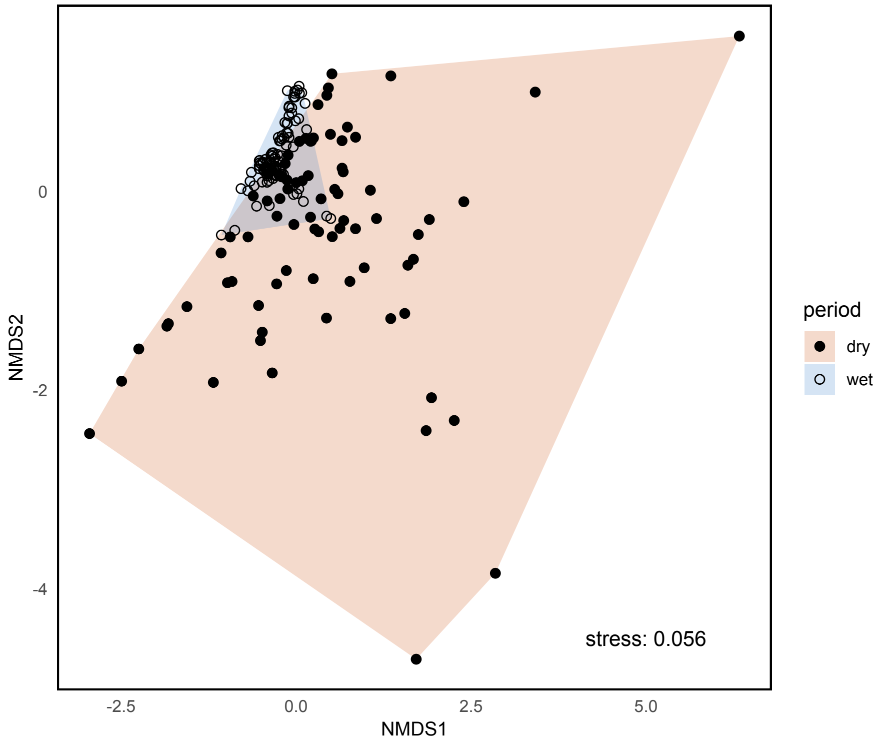


**Figure S7.** NMDS plot showing the changes in environmental conditions (PERMANOVA: F = 31.07, $R^2 = 0.15$, $p = 0.001$; PERMDISP: F = 79.58, $p < 0.001$) in relation to the dry and wet period based on Euclidean distances.

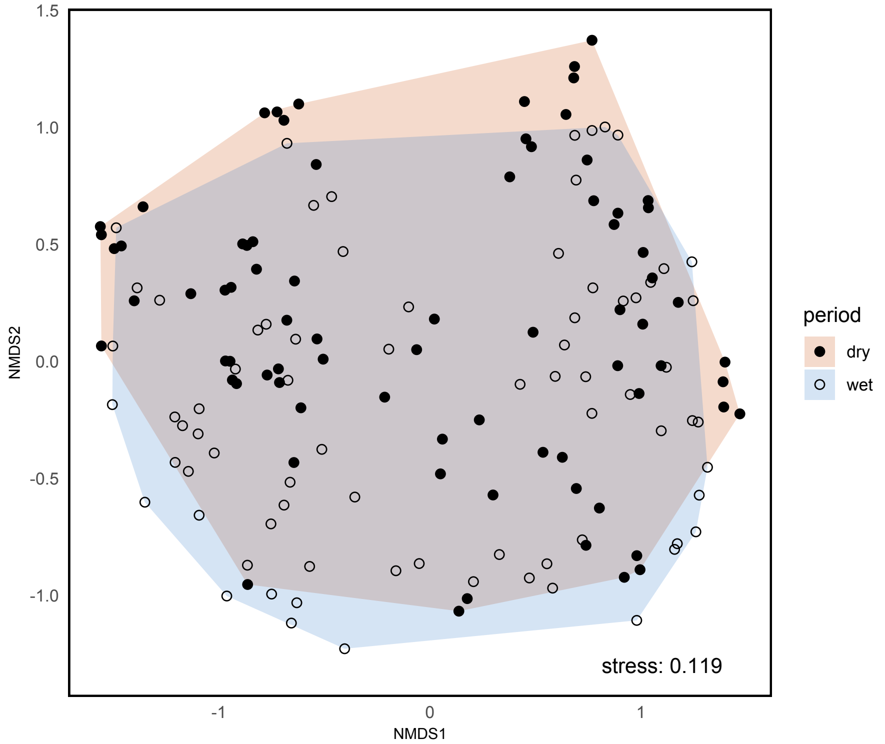


**Figure S8.** NMDS plot showing the changes in bacterial communities (PERMANOVA: F = 3.68, $R^2 = 0.024$, $p = 0.001$; PERMDISP: F = 0.558, $p = 0.456$) in relation to the dry and wet period based on Bray-Curtis dissimilarities.

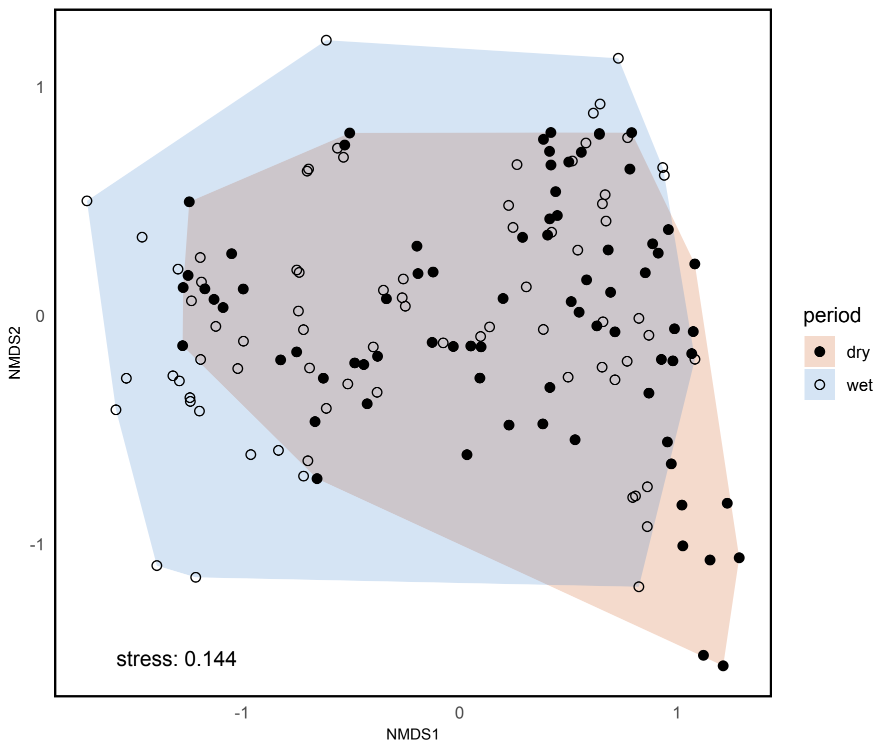


**Figure S9.** NMDS plot showing the changes in microeukaryotic communities (PERMANOVA: F = 5.26, $R^2$ = 0.034, $p$ = 0.001; PERMDISP: F = 1.639, $p$ = 0.202) in relation to the dry and wet period based on Bray-Curtis dissimilarities.

**Table S2.** Separation of environmental conditions, bacterial and microeukaryotic compositions between the dry and wet period in each rock pool based on permutational multivariate analysis of variance (PERMANOVA) using Euclidean distances (for the environmental data) and Bray-Curtis distances (for the Hellinger-transformed OTUs datasets) with 999 permutations. Significant values ($p$ < 0.05) are in bold.

| Rock pool ID | | *Environment* | | | | | | *Bacterial composition* | | | | | | *Microeukaryotic composition* | | | | | |
|---|---|---|---|---|---|---|---|---|---|---|---|---|---|---|---|---|---|---|---|
| | | **df** | **SS** | **MeanSqs** | **F-value** | **$R^2$** | **Pr(>F)** | **df** | **SS** | **MeanSqs** | **F-value** | **$R^2$** | **Pr(>F)** | **df** | **SS** | **MeanSqs** | **F-value** | **$R^2$** | **Pr(>F)** |
| **RP1** | dry/wet | 1 | 10.755 | 10.755 | 7.231 | 0.475 | **0.005** | 1 | 0.251 | 0.251 | 1.204 | 0.131 | 0.275 | 1 | 0.861 | 0.861 | 4.675 | 0.400 | **0.011** |
| | Residuals | 8 | 11.900 | 1.487 | | 0.525 | | 8 | 1.666 | 0.208 | | 0.869 | | 7 | 1.289 | 0.184 | | 0.600 | |
| | Total | 9 | 22.655 | | | 1.000 | | 9 | 1.916 | | | 1.000 | | 8 | 2.150 | | | 1.000 | |
| **RP2** | dry/wet | 1 | 10.031 | 10.031 | 7.680 | 0.490 | **0.011** | 1 | 0.37 | 0.37 | 1.301 | 0.140 | 0.195 | 1 | 0.822 | 0.822 | 5.140 | 0.423 | **0.02** |
| | Residuals | 8 | 10.449 | 1.306 | | 0.510 | | 8 | 2.272 | 0.284 | | 0.860 | | 7 | 1.120 | 0.160 | | 0.577 | |
| | Total | 9 | 20.480 | | | 1.000 | | 9 | 2.642 | | | 1.000 | | 8 | 1.942 | | | 1.000 | |
| **RP3** | dry/wet | 1 | 6.961 | 6.961 | 4.477 | 0.359 | **0.024** | 1 | 0.406 | 0.406 | 1.541 | 0.162 | 0.168 | 1 | 0.521 | 0.521 | 2.404 | 0.231 | **0.041** |
| | Residuals | 8 | 12.440 | 1.555 | | 0.641 | | 8 | 2.105 | 0.263 | | 0.838 | | 8 | 1.733 | 0.217 | | 0.769 | |
| | Total | 9 | 19.401 | | | 1.000 | | 9 | 2.511 | | | 1.000 | | 9 | 2.254 | | | 1.000 | |
| **RP4** | dry/wet | 1 | 7.704 | 7.704 | 4.613 | 0.366 | **0.023** | 1 | 0.585 | 0.585 | 2.219 | 0.241 | **0.043** | 1 | 0.785 | 0.785 | 3.436 | 0.300 | **0.014** |
| | Residuals | 8 | 13.362 | 1.670 | | 0.634 | | 7 | 1.847 | 0.264 | | 0.759 | | 8 | 1.828 | 0.228 | | 0.700 | |
| | Total | 9 | 21.067 | | | 1.000 | | 8 | 2.432 | | | 1.000 | | 9 | 2.613 | | | 1.000 | |
| **RP5** | dry/wet | 1 | 9.395 | 9.395 | 5.305 | 0.399 | **0.007** | 1 | 0.833 | 0.833 | 4.095 | 0.339 | **0.012** | 1 | 0.585 | 0.585 | 2.578 | 0.269 | **0.012** |
| | Residuals | 8 | 14.169 | 1.771 | | 0.601 | | 8 | 1.628 | 0.203 | | 0.661 | | 7 | 1.589 | 0.227 | | 0.731 | |
| | Total | 9 | 23.565 | | | 1.000 | | 9 | 2.461 | | | 1.000 | | 8 | 2.174 | | | 1.000 | |
| **RP7** | dry/wet | 1 | 4.229 | 4.229 | 5.308 | 0.399 | **0.008** | 1 | 0.35 | 0.35 | 1.145 | 0.141 | 0.29 | 1 | 0.949 | 0.949 | 6.321 | 0.441 | **0.008** |
| | Residuals | 8 | 6.374 | 0.797 | | 0.601 | | 7 | 2.139 | 0.306 | | 0.859 | | 8 | 1.201 | 0.150 | | 0.559 | |
| | Total | 9 | 10.604 | | | 1.000 | | 8 | 2.488 | | | 1.000 | | 9 | 2.150 | | | 1.000 | |
| **RP8** | dry/wet | 1 | 12.431 | 12.431 | 11.902 | 0.630 | **0.007** | 1 | 0.467 | 0.467 | 1.641 | 0.190 | 0.053 | 1 | 0.637 | 0.637 | 2.721 | 0.254 | **0.008** |
| | Residuals | 7 | 7.312 | 1.045 | | 0.370 | | 7 | 1.992 | 0.285 | | 0.810 | | 8 | 1.872 | 0.234 | | 0.746 | |
| | Total | 8 | 19.743 | | | 1.000 | | 8 | 2.459 | | | 1.000 | | 9 | 2.509 | | | 1.000 | |
| **RP11** | dry/wet | 1 | 3.706 | 3.706 | 4.929 | 0.413 | **0.012** | 1 | 0.303 | 0.303 | 1.285 | 0.138 | 0.174 | 1 | 0.598 | 0.598 | 4.165 | 0.373 | 0.055 |
| | Residuals | 7 | 5.263 | 0.752 | | 0.587 | | 8 | 1.889 | 0.236 | | 0.862 | | 7 | 1.005 | 0.144 | | 0.627 | |
| | Total | 8 | 8.970 | | | 1.000 | | 9 | 2.192 | | | 1.000 | | 8 | 1.603 | | | 1.000 | |
| **RP12** | dry/wet | 1 | 6.063 | 6.063 | 5.823 | 0.454 | **0.011** | 1 | 0.26 | 0.26 | 0.848 | 0.108 | 0.515 | 1 | 0.863 | 0.863 | 6.750 | 0.458 | **0.017** |
| | Residuals | 7 | 7.289 | 1.041 | | 0.546 | | 7 | 2.146 | 0.307 | | 0.892 | | 8 | 1.023 | 0.128 | | 0.542 | |
| | Total | 8 | 13.351 | | | 1.000 | | 8 | 2.407 | | | 1.000 | | 9 | 1.886 | | | 1.000 | |
| **RP13** | dry/wet | 1 | 8.837 | 8.837 | 9.480 | 0.575 | **0.014** | 1 | 0.412 | 0.412 | 2.16 | 0.236 | **0.018** | 1 | 0.895 | 0.895 | 5.415 | 0.404 | **0.015** |
| | Residuals | 7 | 6.525 | 0.932 | | 0.425 | | 7 | 1.335 | 0.191 | | 0.764 | | 8 | 1.323 | 0.165 | | 0.596 | |
| | Total | 8 | 15.362 | | | 1.000 | | 8 | 1.747 | | | 1.000 | | 9 | 2.218 | | | 1.000 | |
| **RP15** | dry/wet | 1 | 7.988 | 7.988 | 5.703 | 0.449 | **0.006** | 1 | 0.415 | 0.415 | 2.213 | 0.240 | **0.037** | 1 | 1.076 | 1.076 | 8.204 | 0.540 | **0.004** |
| | Residuals | 7 | 9.804 | 1.401 | | 0.551 | | 7 | 1.312 | 0.187 | | 0.760 | | 7 | 0.918 | 0.131 | | 0.460 | |
| | Total | 8 | 17.792 | | | 1.000 | | 8 | 1.727 | | | 1.000 | | 8 | 1.994 | | | 1.000 | |
| **RP16** | dry/wet | 1 | 7.902 | 7.902 | 9.554 | 0.577 | **0.01** | 1 | 0.454 | 0.454 | 1.486 | 0.157 | 0.115 | 1 | 0.602 | 0.602 | 3.248 | 0.289 | **0.008** |
| | Residuals | 7 | 5.790 | 0.827 | | 0.423 | | 8 | 2.443 | 0.305 | | 0.843 | | 8 | 1.482 | 0.185 | | 0.711 | |
| | Total | 8 | 13.692 | | | 1.000 | | 9 | 2.897 | | | 1.000 | | 9 | 2.084 | | | 1.000 | |
| **RP17** | dry/wet | 1 | 4.282 | 4.282 | 2.196 | 0.239 | **0.014** | 1 | 0.47 | 0.47 | 1.638 | 0.170 | 0.091 | 1 | 0.672 | 0.672 | 4.843 | 0.447 | **0.017** |
| | Residuals | 7 | 13.649 | 1.950 | | 0.761 | | 8 | 2.293 | 0.287 | | 0.830 | | 6 | 0.832 | 0.139 | | 0.553 | |
| | Total | 8 | 17.931 | | | 1.000 | | 9 | 2.763 | | | 1.000 | | 7 | 1.504 | | | 1.000 | |
| **RP18** | dry/wet | 1 | 6.223 | 6.223 | 4.835 | 0.409 | **0.005** | 1 | 0.43 | 0.43 | 1.945 | 0.196 | **0.027** | 1 | 0.927 | 0.927 | 5.704 | 0.449 | **0.003** |
| | Residuals | 7 | 9.008 | 1.287 | | 0.591 | | 8 | 1.769 | 0.221 | | 0.804 | | 7 | 1.138 | 0.163 | | 0.551 | |
| | Total | 8 | 15.231 | | | 1.000 | | 9 | 2.2 | | | 1.000 | | 8 | 2.065 | | | 1.000 | |
| **RP19** | dry/wet | 1 | 4.719 | 4.719 | 4.565 | 0.395 | **0.009** | 1 | 0.56 | 0.56 | 3.139 | 0.310 | **0.006** | 1 | 1.066 | 1.066 | 8.329 | 0.510 | **0.005** |
| | Residuals | 7 | 7.236 | 1.034 | | 0.605 | | 7 | 1.25 | 0.179 | | 0.690 | | 8 | 1.024 | 0.128 | | 0.490 | |
| | Total | 8 | 11.955 | | | 1.000 | | 8 | 1.81 | | | 1.000 | | 9 | 2.090 | | | 1.000 | |
| **RP20** | dry/wet | 1 | 5.855 | 5.855 | 3.961 | 0.361 | **0.013** | 1 | 0.608 | 0.608 | 3.049 | 0.303 | **0.018** | 1 | 0.676 | 0.676 | 2.508 | 0.239 | **0.009** |
| | Residuals | 7 | 10.346 | 1.478 | | 0.639 | | 7 | 1.396 | 0.199 | | 0.697 | | 8 | 2.157 | 0.270 | | 0.761 | |
| | Total | 8 | 16.201 | | | 1.000 | | 8 | 2.004 | | | 1.000 | | 9 | 2.833 | | | 1.000 | |

**Table S3.** Separation of environmental conditions, bacterial and microeukaryotic compositions between the dry and wet period in each rock pool based on multivariate homogeneity of group dispersions (PERMDISP) using Euclidean distances (for the environmental data) and Bray-Curtis distances (for the Hellinger-transformed OTUs datasets). Significant values ($p < 0.05$) are in bold.

| **Rock pool ID** | | *Environment* | | | | | *Bacterial composition* | | | | | *Microeukaryotic composition* | | | | |
|---|---|---|---|---|---|---|---|---|---|---|---|---|---|---|---|---|
| | | **df** | **SS** | **MeanSqs** | **F-value** | **Pr(>F)** | **df** | **SS** | **MeanSqs** | **F-value** | **Pr(>F)** | **df** | **SS** | **MeanSqs** | **F-value** | **Pr(>F)** |
| **RP1** | dry/wet | 1 | 0.069 | 0.069 | 0.077 | 0.789 | 1 | 0.004 | 0.004 | 0.167 | 0.694 | 1 | 0.046 | 0.046 | 1.788 | 0.223 |
| | Residuals | 8 | 7.194 | 0.899 | | | 8 | 0.190 | 0.024 | | | 7 | 0.178 | 0.025 | | |
| **RP2** | dry/wet | 1 | 4.434 | 4.434 | 94.698 | **<0.001** | 1 | 0.002 | 0.002 | 0.145 | 0.713 | 1 | 0.078 | 0.078 | 3.528 | 0.102 |
| | Residuals | 8 | 0.375 | 0.047 | | | 8 | 0.116 | 0.014 | | | 7 | 0.155 | 0.022 | | |
| **RP3** | dry/wet | 1 | 1.161 | 1.161 | 3.643 | 0.093 | 1 | 0.011 | 0.011 | 0.389 | 0.550 | 1 | 0.004 | 0.004 | 0.192 | 0.673 |
| | Residuals | 8 | 2.551 | 0.319 | | | 8 | 0.230 | 0.029 | | | 8 | 0.181 | 0.023 | | |
| **RP4** | dry/wet | 1 | 2.331 | 2.331 | 3.577 | 0.095 | 1 | 0.010 | 0.010 | 0.665 | 0.442 | 1 | 0.001 | 0.001 | 0.023 | 0.882 |
| | Residuals | 8 | 5.214 | 0.652 | | | 7 | 0.107 | 0.015 | | | 8 | 0.229 | 0.029 | | |
| **RP5** | dry/wet | 1 | 0.127 | 0.127 | 0.128 | 0.730 | 1 | 0.026 | 0.026 | 4.409 | 0.069 | 1 | 0.001 | 0.001 | 0.181 | 0.683 |
| | Residuals | 8 | 7.986 | 0.998 | | | 8 | 0.046 | 0.006 | | | 7 | 0.044 | 0.006 | | |
| **RP7** | dry/wet | 1 | 1.021 | 1.021 | 3.450 | 0.100 | 1 | 0.001 | 0.001 | 0.017 | 0.900 | 1 | 0.007 | 0.007 | 0.260 | 0.624 |
| | Residuals | 8 | 2.368 | 0.296 | | | 7 | 0.234 | 0.033 | | | 8 | 0.229 | 0.029 | | |
| **RP8** | dry/wet | 1 | 1.555 | 1.555 | 7.378 | **0.030** | 1 | 0.000 | 0.000 | 0.011 | 0.920 | 1 | 0.009 | 0.009 | 0.703 | 0.426 |
| | Residuals | 7 | 1.475 | 0.211 | | | 7 | 0.090 | 0.013 | | | 8 | 0.106 | 0.013 | | |
| **RP11** | dry/wet | 1 | 0.878 | 0.878 | 3.260 | 0.114 | 1 | 0.017 | 0.017 | 0.334 | 0.579 | 1 | 0.076 | 0.076 | 2.633 | 0.149 |
| | Residuals | 7 | 1.885 | 0.269 | | | 8 | 0.418 | 0.052 | | | 7 | 0.201 | 0.029 | | |
| **RP12** | dry/wet | 1 | 0.061 | 0.061 | 0.113 | 0.747 | 1 | 0.001 | 0.001 | 0.014 | 0.908 | 1 | 0.051 | 0.051 | 4.990 | 0.056 |
| | Residuals | 7 | 3.785 | 0.541 | | | 7 | 0.426 | 0.061 | | | 8 | 0.082 | 0.010 | | |
| **RP13** | dry/wet | 1 | 1.538 | 1.538 | 17.674 | **0.004** | 1 | 0.001 | 0.001 | 0.027 | 0.875 | 1 | 0.003 | 0.003 | 0.095 | 0.765 |
| | Residuals | 7 | 0.609 | 0.087 | | | 7 | 0.144 | 0.021 | | | 8 | 0.254 | 0.032 | | |
| **RP15** | dry/wet | 1 | 1.653 | 1.653 | 3.059 | 0.124 | 1 | 0.013 | 0.013 | 1.405 | 0.275 | 1 | 0.006 | 0.006 | 0.840 | 0.390 |
| | Residuals | 7 | 3.781 | 0.540 | | | 7 | 0.063 | 0.009 | | | 7 | 0.054 | 0.008 | | |
| **RP16** | dry/wet | 1 | 1.459 | 1.459 | 7.391 | **0.030** | 1 | 0.001 | 0.001 | 0.040 | 0.846 | 1 | 0.038 | 0.038 | 2.945 | 0.125 |
| | Residuals | 7 | 1.382 | 0.197 | | | 8 | 0.186 | 0.023 | | | 8 | 0.104 | 0.013 | | |
| **RP17** | dry/wet | 1 | 0.167 | 0.167 | 0.139 | 0.720 | 1 | 0.008 | 0.008 | 1.247 | 0.297 | 1 | 0.005 | 0.005 | 0.407 | 0.547 |
| | Residuals | 7 | 8.388 | 1.198 | | | 8 | 0.050 | 0.006 | | | 6 | 0.067 | 0.011 | | |
| **RP18** | dry/wet | 1 | 3.054 | 3.054 | 47.051 | **<0.001** | 1 | 0.035 | 0.035 | 1.793 | 0.217 | 1 | 0.067 | 0.067 | 2.620 | 0.150 |
| | Residuals | 7 | 0.454 | 0.065 | | | 8 | 0.155 | 0.019 | | | 7 | 0.178 | 0.025 | | |
| **RP19** | dry/wet | 1 | 1.777 | 1.777 | 4.377 | 0.075 | 1 | 0.031 | 0.031 | 2.099 | 0.191 | 1 | 0.041 | 0.041 | 2.837 | 0.131 |
| | Residuals | 7 | 2.842 | 0.406 | | | 7 | 0.102 | 0.015 | | | 8 | 0.115 | 0.014 | | |
| **RP20** | dry/wet | 1 | 2.775 | 2.775 | 3.593 | 0.100 | 1 | 0.020 | 0.020 | 0.508 | 0.499 | 1 | 0.014 | 0.014 | 1.507 | 0.255 |
| | Residuals | 7 | 5.408 | 0.773 | | | 7 | 0.275 | 0.039 | | | 8 | 0.075 | 0.009 | | |

*Results of the elements of metacommunity structure (EMS) analyses*

**Table S4.** Results of EMS analysis for bacterial and microeukaryotic communities based on fixed-proportional null models performed separately on matrices ranked based on the first ordination (primary) axis extracted via reciprocal averaging. *Abs*: number of embedded absences, *Turn*: number of replacements. Interpretations follow Presley et al. [1]. Significant results ($p$ < 0.05) are in bold.

**Bacterioplankton**

| | Sampling | Percent | Coherence | | | | | Turnover | | | | | Clumping | | | Metacommunity |
|---|---|---|---|---|---|---|---|---|---|---|---|---|---|---|---|---|
| | **dates** | **interia** | **Abs** | **z** | **p** | **sim.Mean** | **sim.Var** | **Turn** | **z** | **p** | **sim.Mean** | **sim.Var** | **index** | **p** | **df** | **type** |
| Primary axis | 2015-08-14 | 11.6% | 21705 | -0.220 | 0.825 | 21529.6 | 795.5 | 7147070 | -6.564 | **<0.0001** | 6566729.5 | 88408.9 | 1.156 | **<0.0001** | 12 | Random |
| | 2015-08-18 | 12.1% | 23389 | -1.555 | 0.120 | 22061.2 | 853.8 | 8101347 | -1.588 | 0.112 | 7936575.0 | 103784.0 | 1.097 | **<0.0001** | 11 | Random |
| | 2015-08-22 | 10.9% | 33565 | -2.359 | **0.018** | 30909.2 | 1125.7 | 13866336 | -4.329 | **<0.0001** | 13141599.3 | 167417.5 | 1.303 | **<0.0001** | 13 | Checkerboards |
| | 2015-08-26 | 11.1% | 27107 | 1.787 | 0.074 | 28800.8 | 947.9 | 12817243 | -10.033 | **<0.0001** | 11492551.9 | 132036.7 | 1.156 | **<0.0001** | 13 | Random |
| | 2015-08-30 | 10.5% | 28358 | 1.244 | 0.213 | 29656.5 | 1043.8 | 13779751 | -12.706 | **<0.0001** | 11959169.3 | 143282.6 | 1.281 | **<0.0001** | 13 | Random |
| | 2015-09-03 | 12.1% | 33743 | -0.075 | 0.940 | 33657.4 | 1136.3 | 20594763 | -9.206 | **<0.0001** | 18575990.0 | 219277.5 | 1.109 | **<0.0001** | 13 | Random |
| | 2015-09-07 | 11.6% | 27946 | -0.493 | 0.622 | 27477.0 | 951.4 | 12822665 | -9.318 | **<0.0001** | 11527035.1 | 139050.0 | 1.135 | **<0.0001** | 12 | Random |
| | 2015-09-11 | 12.7% | 21750 | 3.976 | **<0.0001** | 25194.9 | 866.3 | 11178647 | -17.021 | **<0.0001** | 9238071.1 | 114009.0 | 1.067 | **<0.0001** | 12 | Nested - Clumped species loss |
| | 2015-09-15 | 13.4% | 22805 | 1.483 | 0.138 | 24097.7 | 871.7 | 9307139 | -12.374 | **<0.0001** | 8131338.5 | 95020.3 | 1.163 | **<0.0001** | 12 | Random |
| | 2015-09-19 | 12.3% | 17095 | 0.718 | 0.473 | 17635.0 | 752.4 | 4750609 | -5.066 | **<0.0001** | 4416531.7 | 65951.5 | 1.157 | **<0.0001** | 11 | Random |

**Microeukaryotes**

| | Sampling | Percent | Coherence | | | | | Turnover | | | | | Clumping | | | Metacommunity |
|---|---|---|---|---|---|---|---|---|---|---|---|---|---|---|---|---|
| | **dates** | **interia** | **Abs** | **z** | **p** | **sim.Mean** | **sim.Var** | **Turn** | **z** | **p** | **sim.Mean** | **sim.Var** | **index** | **p** | **df** | **type** |
| Primary axis | 2015-08-14 | 12.6% | 4593 | -1.823 | 0.068 | 4213.8 | 208.0 | 346150 | 0.163 | 0.870 | 347921.7 | 10854.6 | 1.187 | **<0.0001** | 10 | Random |
| | 2015-08-18 | 11.1% | 7951 | **-2.484** | **0.013** | 7264.8 | 276.2 | 921033 | 2.147 | **0.032** | 989418.8 | 31858.2 | 1.128 | **<0.0001** | 13 | Checkerboards |
| | 2015-08-22 | 13.6% | 8291 | -1.372 | 0.170 | 7850.7 | 321.0 | 1159339 | 2.200 | **0.028** | 1232054.0 | 33050.4 | 1.138 | **<0.0001** | 13 | Random |
| | 2015-08-26 | 12.0% | 6803 | -1.280 | 0.200 | 6455.0 | 271.8 | 766156 | 2.940 | **0.003** | 836760.9 | 24013.7 | 1.124 | **<0.0001** | 12 | Random |
| | 2015-08-30 | 11.8% | 8068 | -1.485 | 0.138 | 7642.9 | 286.2 | 1171495 | -1.833 | 0.067 | 1117742.8 | 29318.8 | 1.064 | **<0.0001** | 13 | Random |
| | 2015-09-03 | 12.7% | 9842 | **-2.440** | **0.015** | 8951.0 | 365.2 | 1618172 | 3.406 | **0.001** | 1769959.3 | 44569.6 | 1.140 | **<0.0001** | 12 | Checkerboards |
| | 2015-09-07 | 13.4% | 7779 | 1.125 | 0.260 | 8159.6 | 338.2 | 1437851 | -8.138 | **0.000** | 1193279.8 | 30051.5 | 1.195 | **<0.0001** | 12 | Random |
| | 2015-09-11 | 10.6% | 8767 | -0.899 | 0.369 | 8473.5 | 326.4 | 1440551 | -7.356 | **0.000** | 1213772.3 | 30828.4 | 1.129 | **<0.0001** | 13 | Random |
| | 2015-09-15 | 11.5% | 7250 | -0.237 | 0.813 | 7183.4 | 281.1 | 890580 | -3.938 | **0.000** | 811100.0 | 20181.0 | 1.100 | **<0.0001** | 12 | Random |
| | 2015-09-19 | 11.5% | 7001 | -0.486 | 0.627 | 6857.7 | 294.6 | 860170 | 0.016 | 0.987 | 860478.6 | 19263.6 | 1.092 | **<0.0001** | 12 | Random |

*Data processing*

Raw sequence data were analysed and sequences clustered into OTUs (97% similarity) according to the UPARSE pipeline [2], as described on the BILS website for using UPARSE on the UPPMAX cluster (available with all scripts at: https://wiki.bils.se/wiki/Running_the_Uparse_pipeline_at_the_UPPMAX_cluster). We further assigned taxonomy to the representative 16S and 18S OTUs using the SSU Ref NR 99 v119 SILVA database [3]. Chloroplast OTUs and OTUs that were unassigned taxonomically or represented by less than 10 sequences were discarded. To generate an equal number of sequences per sample the OTU table was subsampled based on the sample with the lowest number of reads (4 940 reads/sample in the bacterial dataset, 3 429 reads/sample in the microeukaryotic dataset). A total of 7 767 202 quality controlled 16S rRNA sequencing reads were acquired. After OTU clustering (sample concatenation, dereplication and sorting) 5 291 921 sequences were taxonomically assigned (with a 95% sequence identity threshold). After removal of non-bacterial sequences and subsampling, a total of 4 587 OTUs were retained for the bacterial communities. In the 18S rRNA dataset, 6 349 385 good-quality reads were gained and 3 776 690 sequences were taxonomically assigned (with 95% sequence identity threshold) after OTU clustering. After removal of non-eukaryotic OTUs and subsampling, the microeukaryotic dataset consisted of 1 336 OTUs. The taxonomic distribution of reads for both datasets was visualized with Krona (http://sourceforge.net/projects/krona).
To construct phylogenetic trees, first, the representative sequences of each OTU were aligned to the Greengenes core reference alignment [4] in the case of bacteria, and to the SILVA 128 core reference alignment [3] in the case of microeukaryotes using PyNAST [5] in MacQIIME v1.9.1 [6]. To remove gaps and highly variable regions, the alignments were filtered using a Lane mask [7] and were then used to construct approximate maximum-likelihood phylogenetic trees with FastTree v2.1.3.

*Detailed description of the method of elements of metacommunity structure (EMS)*

We examined coherence where statistical significance of the number of embedded absences was determined with z-tests resulting in z-value of coherence. Non-significant z-values (no coherence) suggest randomly structured assemblages. z-values that are significantly lower than in the null distribution (negative coherence) suggest checkerboard patterns due to competitively structured assemblage. Finally, z-values that are significantly higher compared to the null model distribution (positive coherence) indicate that assemblages are structured along environmental gradients, either individualistically or by coherent units of species that respond similarly to the changes in the environment. In the next step we therefore examined species turnover based on z-tests to depict the character of these gradual changes. If turnover was significantly higher than expected by chance, this is indicative of species sorting along environmental gradients and boundary clumping based on Morisita's index was used to describe whether individualistic (Gleasonian pattern, replacement of individual species along the gradient) or synchronous (Clementsian pattern, replacement of groups of species along the gradient) species turnover was important. If turnover is significantly lower than expected by chance, the metacommunity had nested subsets in which species in less diverse communities are subsets of those in more diverse communities [8].
